# Synergistic interactions among *Burkholderia cepacia* complex (Bcc)-targeting phages reveal a novel therapeutic role for lysogenization-capable (LC) phages

**DOI:** 10.1101/2022.10.26.513969

**Authors:** Philip Lauman, Jonathan J. Dennis

## Abstract

Antimicrobial resistance is an imminent danger to global public health and threatens virtually all aspects of modern medicine. Particularly concerning are the species of the *Burkholderia cepacia* complex (Bcc), which cause life-threatening respiratory infections among patients who are immunocompromised or afflicted with cystic fibrosis, and are notoriously resistant to antibiotics. One promising alternative being explored to combat Bcc infections is phage therapy (PT) - the use of phages to treat bacterial infections. Unfortunately, the utility of PT against many pathogenic species, including the Bcc, is limited by the prevailing paradigm of PT: that only obligately lytic phages, which are rare, should be used therapeutically - due to the conviction that so-called ‘lysogenic’ phages do not reliably clear bacteria and instead form lysogens to which they may transfer antimicrobial resistance or virulence factors. In this study, we argue that the tendency of a lysogenization-capable (LC) phage to form stable lysogens is not predicated exclusively on its ability to do so, and that this property, along with the therapeutic suitability of the phage, must be evaluated on a case-by-case basis. Concordantly we developed several novel metrics - Efficiency of Phage Activity (EPA), Growth Reduction Coefficient (GRC), and Lysogenization Frequency (*f*_*(lys)*_) and used them to evaluate eight phages targeting members of the Bcc. We found that while these parameters vary considerably among Bcc phages, a strong inverse correlation exists between lysogen-formation and antibacterial activity, indicating that certain LC phages may be highly efficacious on their own. Moreover, we show that many LC Bcc phages interact synergistically with other phages in the first reported instance of mathematically defined polyphage synergy, and that these interactions result in the eradication of *in-vitro* bacterial growth. Together, these findings reveal a novel therapeutic role for LC phages, and challenge the current paradigm of PT.

**IMPORTANCE:** The spread of antimicrobial resistance is an imminent threat to public health around the world. Particularly concerning are the species of the *Burkholderia cepacia* complex (Bcc), which cause life-threatening respiratory infections and are notoriously resistant to antibiotics. Phage therapy (PT) is a promising alternative being explored to combat Bcc infections and antimicrobial resistance in general, but the utility of PT against many pathogenic species, including the Bcc, is restricted by the currently prevailing paradigm of exclusively using rare obligately lytic phages - due to the perception that ‘lysogenic’ phages are therapeutically unsuitable. Our findings show that many lysogenization-capable (LC) phages exhibit powerful *in vitro* antibacterial activity both alone and through mathematically defined synergistic interactions with other phages, demonstrating a novel therapeutic role for LC phages and therefore challenging the currently prevailing paradigm of PT.

## INTRODUCTION

The increasing prevalence and global distribution of multidrug resistant (MDR) bacterial pathogens is an imminent menace to public health and threatens virtually all aspects of modern medicine. Antibiotic resistance is a major cause of mortality in both developed and developing countries worldwide, and it is currently predicted that drug resistance will cause in excess of 10 million premature deaths per year by 2050, and cost an approximated USD 100 trillion in damages (1–3). Discovery of new antibiotics with novel cellular targets has failed to keep pace with the evolution of bacterial resistance, which has now emerged even against antibiotics of last resort, meaning that novel modes of treatment are desperately needed to combat resistant pathogens (4, 5).

Among the most problematic of these pathogens are the members of the *Burkholderia cepacia* complex (Bcc), a group of over twenty notoriously drug-resistant species which cause severe disease in patients who are immunocompromised or afflicted with certain diseases – notably chronic granulomatous disease (CGD) and cystic fibrosis (CF; 6–8). Although Bcc members account for a relatively small proportion of pulmonary infections in both CF and CGD populations, infections by these organisms are remarkably aggressive and have the highest mortality rates in both of these patient groups. In many cases, high fatality rates are due to a severe disease progression termed “*cepacia* syndrome”, which contrasts with the clinical progressions of infections caused by other CF and CGD pathogens and is characterized by high fever, severe progressive respiratory failure, decline in leukocyte and erythrocyte levels, necrosis, bacteremia, and sepsis – which result in rapid death (8–12). Moreover, at least five Bcc species – *B. cepacia, Burkholderia cenocepacia, Burkholderia multivorans, Burkholderia dolosa*, and *Burkholderia contaminans* – are capable of spreading via aerosol droplets, meaning these pathogens can disseminate readily, through direct and indirect contact, among susceptible patients in hospital settings. As a result, Bcc-positive patients are required to follow stringent infection control practices and are isolated from other patients in CF clinics – measures that have serious psychosocial ramifications which can potentially exacerbate patient illness (13–15). Finally, Bcc species possess a large repertoire of both innate and acquired antimicrobial resistance mechanisms, including upregulated efflux pumps, porins, and *β*-lactamases, outer membrane modifications resulting in decreased permeability, as well as mutations in key antibiotic targets such as dihydrofolate reductase and penicillin binding proteins. Collectively, these mechanisms confer resistance to most antibiotics, including *β*-lactams, aminoglycosides, antimicrobial peptides, polymixins, most quinolones and tetracyclines, chloramphenicol, and trimethoprim (7, 8). Even the most effective anti-Bcc antibiotics – ceftazidime, meropenem, and minocycline – have been reported to inhibit only 23-38% of clinical isolates in the United States, meaning that conventional antibiotic treatments are largely ineffective and alternative approaches to treating Bcc infections are therefore urgently required (12, 16, 17).

One alternative strategy being explored to combat Bcc infections, and antimicrobial resistance in general, is phage therapy – the clinical administration of bacteriophages (or phages) to treat bacterial infections. Phages are bacterial viruses which readily infect and destroy bacterial cells in order to replicate themselves and are environmentally abundant, outnumbering the bacterial population by approximately tenfold. Importantly, phage predation produces an estimated 10^23^ infections per second and destroys roughly 50% of the global bacterial population every 48 h, meaning that phage therapy is merely the application of a naturally occurring antibacterial agent to a human problem (8, 18–20). In nature, the tailed phages of the order *Caudovirales* – which are most relevant to phage therapy – most commonly employ two evolutionarily complementary replication cycles, termed the lytic and lysogenic cycles. In the lytic cycle, phages begin the sequential process of genome replication, protein expression, virion assembly, and cell lysis immediately upon entry into a host cell, resulting in relatively rapid destruction of the cell and, by extension, the host population. In the complementary lysogenic cycle, the phage genome is incorporated into that of the host and replicates passively with the host cell until conditions trigger re-induction to the lytic cycle. Lysis of the host cell is therefore the ultimate result of both of these replication cycles, but is generally achieved more slowly through the lysogenic cycle (8, 21, 22).

In keeping with the terminology used to describe the replication cycles of therapeutically relevant *Caudovirales*, these phages have canonically been dichotomously categorized as either “lytic” or “lysogenic”, designations with which the terms “virulent” and “temperate”, respectively, have often been used interchangeably. The current use of this terminology is quite problematic, however, since so-called “lysogenic” phages can still produce lysis and can be virulent in the sense that they kill bacteria or reduce bacterial growth. In recent years, some researchers have begun utilizing the term obligately lytic (OL; also called strictly or professionally lytic) to more accurately describe phages that are genetically incapable of forming lysogens, while the term “virulent” has been mathematically defined by Storms et al. as highly capable of reducing bacterial growth during the logarithmic phase (23–25). Nevertheless, similar elaboration has not been achieved thus far on the terms “lysogenic” or “temperate”. Since the tendency to form lysogens is not predicated solely on the mere ability to do so, and depends on host and environmental factors as well (26, 27), it is inaccurate to say that all phages which are genetically capable of forming lysogens do so to any meaningful degree under all possible environmental conditions, including conditions which are therapeutically relevant. Concordantly, we propose that these phages should be described as lysogenization-capable (LC), rather than lysogenic, while the latter term should be reserved for description of the lysogenic replication cycle. Rather than being dichotomously categorized as either “lytic” or “lysogenic”, individual phages should be understood as occupying points on a spectrum in terms of their tendency to form lysogens under a particular set of environmental conditions, with OL phages occupying one of the outer boundaries of this spectrum. As well, the term “temperate” implicitly suggests a lack of lytic activity and virulence – which is not always the case among lysogenization-capable phages (28–31), and this term should therefore be used as an antonym to “virulent”, rather than as a synonym for lysogenization-capable.

Although the antibacterial effects of such LC phages have been explored, both *in vitro* and *in vivo*, in a handful of studies over the past decade (24, 29, 31–34), the prevailing paradigm within phage therapy is that only OL phages are therapeutically suitable, while LC phages are therapeutically suboptimal. This preference is due to the perception that if they form lysogens, LC phages (1) may not effectively eliminate the targeted bacterial population and (2) may provide virulence or antimicrobial resistance factors to target bacteria (8, 23, 24, 27, 35). As previously discussed, however, the mere ability of a phage to produce lysogens is not the sole predictor of whether the phage will do so under therapeutically relevant conditions. Furthermore, even if such phages do form lysogens – they may not have virulence or antimicrobial resistance factors to transfer. Finally, even if certain phages form lysogens under therapeutically relevant conditions, this only implies that they cannot be used on their own through monophage therapy. Combining these phages with OL phages or LC phages which do not form lysogens under therapeutic conditions could still form therapeutically efficacious polyphage cocktails. As a result, LC phages should not be dismissed outright based on their genetic properties, but instead ought to be evaluated for their tendency to form stable lysogens, and their ability to reduce bacterial growth – both alone and in combination with other phages – on a case-by-case basis.

The aforementioned shortcomings in the current paradigm of phage therapy might be harmless if OL phages were abundant, but these phages are in fact challenging to find for many therapeutically problematic bacterial species, while LC phages appear to be ubiquitous (24, 27, 36). Among the Bcc, for instance, only five OL phages have ever been identified, and only one of these has been characterized (8, 37, 38). This is unsurprising from an evolutionary perspective, since the possession of a lysogenic cassette is evolutionarily beneficial for the long-term stability of phage populations (19, 27, 39, 40). It is reasonable, then, to expect that OL phages may be the minority of all existing phages, and the current paradigm of only using such phages is therefore not only illogical but also impractical as it may restrict the range of bacterial species against which phage therapy can be utilized.

It is thus clear that a case-by-case evaluation of LC phages is necessary to validate their therapeutic suitability. To further this goal for phages targeting the Bcc – a group of pathogenic species against which polyphage cocktail development is desperately required but has never previously been described in literature – we developed novel metrics to gauge the tendency of Bcc phages to form stable lysogens, as well the capacity of these phages to reduce bacterial growth *in vitro*, both alone and in combination with another phage. We then investigated these properties for eight previously characterized Bcc phages, seven LC and one OL, on seventeen variably susceptible strains of the Bcc. Here we present the trends identified for these variables, as well as the relationships between them, and demonstrate that at least some LC phages have enormous therapeutic potential – not only via monophage treatment – but also in combinations in which they synergize with other phages to effectively abolish bacterial growth.

## METHODS

### Organisms & Media

Seventeen strains belonging to the Bcc were examined as part of this study, including 9 *B. cenocepacia*, 2 *Burkholderia ambifaria*, 2 *B. multivorans*, 1 *Burkholderia stabilis*, and 3 *B. dolosa*. 15 of these strains were clinical isolates from CF patients in several countries including Canada (6), USA (5), Australia (2), UK (1), and Belgium (1), while two strains were environmental isolates from rhizosphere samples (**Supplementary Table 1**). Virtually all experiments in this study were conducted using half-strength Luria-Bertani broth (½ LB (41); 5g/L Bacto tryptone, 2.5 g/L yeast extract, 2.5 g/L NaCl) or ½ LB 1.5% (w/v) agar plates, while 20% glycerol full-strength LB was used for long-term storage of bacterial strains at -80°C. Bacteria were incubated statically on ½ LB 1.5% (w/v) agar plates at 30°C for 48hrs or until colonies appeared, and single colonies were cultured in 5mL ½ LB broth at 30°C and at 225 rpm in a gyratory shaker for precisely 18hrs. At 18hrs, bacterial cultures were plated for viable CFUs and their optical density at 600nm (OD_600_) were measured using a Nanodrop spectrophotometer. OD_600_-CFU equivalencies for this timepoint were then constructed for every bacterial strain, and were used to approximate viable CFUs in 18hr cultures. Bacterial cultures were serially diluted in fresh ½ LB broth to a starting inoculum of 10^5^ or 10^3^ CFU (see **Supplementary Table 1**), which roughly corresponds to the LD_50_ of most Bcc strains in *Galleria mellonella* greater wax moth larvae (42).

Eight Bcc phages, previously isolated and characterized by members of the Dennis Laboratory (8, 12, 38, 43–46), including seven LC phages AH2, DC1, KL1, KS4M, KS5, KS9, KS14 and OL phage JG068, were examined as part of this study (**Supplementary Table 2**). All phages were stored at 4°C in Suspension Medium (SM (46); 5.8g/L NaCl, 2g/L MgSO_4_, 50mL/L 1M Tris-HCl) until required, and maintenance of high phage titre was routinely verified. Determination of phage titre was conducted using the double-layer agar approach (32), and this technique (with collection of lysate) was performed repeatedly until sufficiently high phage titres (10^9^-10^10^ PFU/mL) were obtained. Phages DC1 and KS14 were amplified only on *B. cenocepacia* C6433 and phages KS9 and JG068 only on *B. cenocepacia* K56-2, while phages AH2, KL1, KS4M and KS5 were amplified on both of these strains, giving rise to variants with slightly altered host ranges (not shown), but only the more effective (on relevant strains) variants of these phages were used in this study.

### Efficiency of Phage Activity (EPA) Assay

This method is procedurally similar to the previously described Efficiency of Plating (EOP) Assay (47) in which a confluent lawn of the target bacterium is poured onto a ½ LB 1.5% (w/v) agar plate using the double-agar method described previously (32). Phages are serially diluted in SM in order of magnitude steps from 10^10^ to 10^3^ PFU/mL, and 0.005mL of each concentration is then spotted in triplicate onto the bacterial lawn, meaning the lowest concentration at which a spot would contain at least one phage is 10^3^ PFU/mL, which is therefore the lowest concentration at which evidence of phage activity is theoretically possible. Plates are incubated statically at 30°C for 18hrs and are then inspected visually. The lowest concentration at which evidence of phage activity (plaques, mottling, turbid clearance) is visible is then compared to the lowest concentration at which such evidence is theoretically possible using **Equation 1** (see *results*) to obtain the EPA score. EPA scores are unitless and range from a maximum of 0 to a minimum of - 7, while EPA < -7 indicates lack of sensitivity to phage activity.

### Planktonic Killing Assay (PKA)

All experiments were conducted using flat bottom 24-well plates (Corning, USA). 0.1mL of a 10^6^ CFU/mL diluted culture was added to each relevant well and was mixed with 0.1mL of phage stock at order-of-magnitude concentrations ranging from 10^5^ to 10^10^ PFU/mL, thereby generating an MOI range of 10^−1^ to 10^4^. Slightly different values were used for *B. cenocepacia* K56-2 (see **Supplementary Table 1**) but the same MOI range was maintained. Assays with two-phage combinations were performed using the same protocol, but only at the maximum available MOI, either 10^3^ or 10^4^. Bacteria and phage were permitted to mix for 10 min at room temperature and were subsequently covered with 2mL fresh ½ LB broth. Positive control wells contained 0.1mL of bacteria mixed with 0.1mL SM, covered with 2mL ½ LB, while negative control wells received only 0.1mL SM and were covered with 2.1mL ½ LB to maintain identical volume. Plates were incubated in either a BioTek Epoch 2 microplate spectrophotometer or BioTek Cytation 5 Multi-Mode Reader at 30°C with continuous shaking at 237rpm for 48hrs, and OD_600_ readings were collected every hour. At least three biological replicates were conducted at each MOI for each phage and two-phage combination, and data from biological replicates was then averaged to produce a single growth curve at each MOI. Generated growth curves were analyzed using the Growth Reduction Coefficient (GRC) approach, which uses the area under the growth curve (AUC) to provide an estimate of the ability of the phage (or cocktail) to reduce bacterial growth (see results).

### Lysogenization Frequency (*f*_(*lys*)_) Assays

*In-vitro f*_(*lys*)_ experiments were performed using flat bottom 24-well plates (Corning, USA), with the same bacterial inoculum, phage concentration and MOI ranges, and general procedures as described for PKAs, but four biological replicates were conducted for each phage-host pair and plates were incubated at either 30°C or 37°C at 225rpm in a gyratory shaker for precisely 48hrs. At 48hrs, 1mL of surviving cells were centrifuged for 5 min at 12 000rpm to remove phage and were washed with 0.5mL of fresh ½ LB. This procedure was repeated thrice to remove phage contamination and surviving cells were plated for single colony isolation. After 48hrs of static growth at 30°C, 15 colonies were randomly selected from each of four plates (biological replicates), for a total of 60 colonies per phage. Colonies were PCR-screened for lysogeny status and average *f*_(*lys*)_ values for the four replicates were calculated using **Equation 9** (see results). *In-vitro* assays with KS14 were conducted in M9 Minimal Medium (M9 MM (48) ; 200mL/L 5x M9 Salts, 2g/L MgSO_4_, 0.1g/L CaCl_2_, 4g/L glucose) and Artificial Cystic Fibrosis Sputum Medium (ACFSM ; 49), in addition to ½ LB, while assays with all other phages were conducted only in ½ LB. *In vivo f*_(*lys*)_ experiments were performed only with phage KS14 in the haemolymph of *Galleria mellonella* greater wax moth larvae. For each KS14-host pair, 12 larvae (3 technical replicates x 4 biological replicates) were injected sequentially with 10^5^ CFU and 10^8^ PFU of KS14 (MOI: 10^3^), with a 2hr interval between injections, as previously described (42), and larvae were incubated at 30°C in the dark for 48hrs after injection. After 48hrs, larvae were euthanized via decapitation and haemolymph was extracted. The haemolymph of technical replicate larvae was pooled to form four biological replicate stocks, from which survivor colonies were isolated and PCR screened for lysogeny status as described above.

### Statistical Analysis

In all figures and tables, data points for GRC and *f*_(*lys*)_ represent the mean values of at least three and four biological replicates, respectively. Error bars represent standard error of the mean (SEM) of all biological replicates, and errors reported for 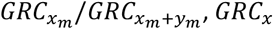, and 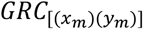 values (see **equations 3, 6**, and **11**, respectively, in results) are the summation of errors reported for their components. GraphPad Prism 9.3.1 (California, USA) was used to compute the areas under all bacterial growth curves using the trapezoid approximation method, as well as to conduct all statistical analyses including linear regression between variables, and Student’s T-tests (with Welch’s correction for unequal variances) for comparisons between the GRCs of single phages and two-phage cocktails. *p*<0.05 was considered statistically significant.

## RESULTS

### Efficiency of Phage Activity (EPA) is a novel quantitative approach for screening phages on solid medium

Efficiency of Plaquing (EOP; also called bacteriophage Efficiency of Plating) is a widely used metric for determining the host range of a phage, and the efficiency with which it is able to infect a given host bacterium on solid medium (47, 50, 51). While obviously useful, the fact that this approach requires the counting of discrete plaques means it cannot be used when working with phages which do not form plaques and instead produce turbid clearing or mottling – which is sometimes indicative of non-productive activity such as abortive infection or lysis from without but can also indicate low levels of productive infection (52, 53). Since many Bcc phages produce such non-plaque evidence phage activity on certain hosts, we devised an alternative approach – designated Efficiency of Phage Activity (EPA) – which uses all possible evidence of infection (including mottling, turbid clearing, and plaquing) rather than relying exclusively on plaques. In this approach, the EPA of a phage *x* on a given host strain *h* is computed as the logarithm of the quotient of the lowest order-of-magnitude concentration at which evidence of infection is *theoretically* possible, [*P*_*x*_]_*L*(*T*)_, and the lowest order-of-magnitude concentration at which evidence of infection is *actually* seen, [*P*_*x*_]_*L*(*A*)_, as shown in **equation 1**:

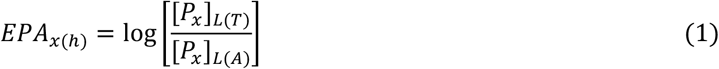

EPA thus provides an order-of-magnitude estimate of a given phage’s activity on a particular bacterial strain on solid medium, even if it is unable to form discrete plaques. This allows direct comparison of the activities of different phages against particular strains, and thus serves as a starting point for evaluating which phage-host pairs could be explored for therapeutic applications. Using this modified approach, we quantified the EPA of our eight Bcc phages on a panel of seventeen strains belonging to five species of the Bcc (**Supplementary Figure 1**). We found that the EPA varies substantially by host strain for most tested Bcc phages with the exception of the OL phage JG068 – which has a high EPA on all tested host strains. Interestingly, many phages have host ranges which differ from those reported in previous literature (8), suggesting modified host affinities possibly resulting from adaptation in the lab. Since this approach is intended to serve as a preliminary screen to identify phages with high levels of activity against particular strains, we utilized an arbitrary EPA threshold of EPA ≥ -5 and pursued subsequent experiments with phage-host pairs which satisfied this requirement, while those that did not meet the requirement were excluded from further studies.

### The Growth Reduction Coefficient (GRC), a novel metric for quantification of a phage’s ability to reduce bacterial growth, reveals enormous diversity in the antibacterial capabilities of Bcc phages

Since *in-vivo* infection dynamics are, in most cases, better represented by infections in liquid medium rather than solid medium (25, 51, 54, 55), we sought to evaluate the *in-vitro* antibacterial abilities of Bcc phages through planktonic killing assays (PKA). Within the field of phage therapy, studies in which qualitative assessments are made regarding the effects of phage administration on bacterial growth curves are abundant (32, 56, 57), but a method to interpret these effects quantitatively was previously unavailable. In a recent study, Storms et al. made a major contribution to this field by developing a novel variable termed the Virulence Index (VI), which quantifies the ability of a phage to reduce bacterial growth during the logarithmic growth phase (25). Here we present a therapeutically relevant variant of this metric – designated the Growth Reduction Coefficient (GRC) – which quantifies the ability of a phage to reduce bacterial growth across the entirety of a therapeutically relevant period of time – which may vary depending on the typical clinical course of infection by the particular bacterial species being investigated. Because GRC is measured across such an extensive time period, rather than the logarithmic growth phase of the bacterium, it can detect whether a phage is capable of reducing bacterial growth after the stationary phase has been reached, or of preventing resistant outgrowths which may appear at a later time – an advantage which is particularly relevant when investigating the antibacterial potential of phage-phage or phage-antibiotic combinations. In presenting the VI, Storms et al. found that virulence varies substantially with the growth medium and temperature being utilized (25), so it is logical to assume that similar variability would be found when using the GRC metric. In this study, however, we focus on a particular set of infection conditions which were used in previous *in-vitro* work with Bcc phages.

In this approach, a target bacterial strain is grown in the presence and absence of a phage *x* at various multiplicities of infection (MOI, *m*), and we utilize the definite integral of the generated *OD* _600_ vs *t* growth curve (see left panels in **Figure 1**) to define the related area under the curve, 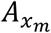, as shown in **equation 2**:

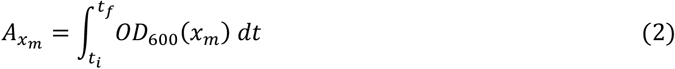

The Growth Reduction Coefficient of phage *x* at MOI *m*, 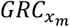 (designated the *local* GRC at MOI *m*, in keeping with the terminology devised by Storms et al.), is then defined through comparison with the area under the growth curve of the untreated bacterial strain, *A*_0_, as shown in **equation 3:**

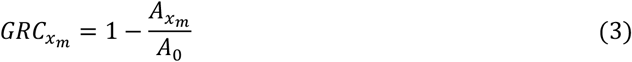

The GRC is thus expressed as a unitless value which can range up to 1, which represents complete abolition of bacterial growth, while positive values near zero reflect a poor ability to reduce bacterial growth and negative values conversely suggest the tendency to improve bacterial growth. 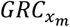 values can thus be calculated for a range of relevant MOIs, and can subsequently be used to construct a 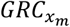 vs *log*_10_*MOI* curve (see right panels in **Figure 1**). The area under this curve, designated *A*_*x*_, is defined using **equation 4**:

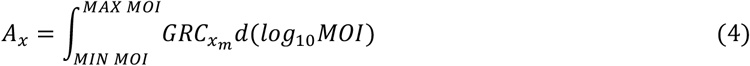

The maximum theoretical area under this curve, designated *A*_*MAX*_, is then computed using the number of MOI conditions utilized, *n*_*m*_, as shown in **equation 5**:

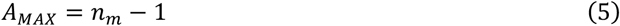

The GRC of phage *x* across all tested MOIs, *GRC*_*x*_ (designated the *global* GRC of phage *x*), is then defined as the quotient of *A*_*x*_ and *A*_*MAX*_, as shown in **equation 6**:

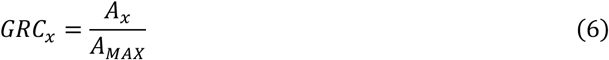

*GRC*_*x*_ and 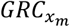 values can subsequently be compared between the same phage on different hosts, or different phages on the same host, to quantify the potential therapeutic suitability of various phages.

GRC experiments of this type were conducted for all Bcc phage-host pairs for which an EPA ≥ -5 was obtained (*n*=37; representative data in **Figure 1**; all data in **Supplementary Figure 2**). When investigating Bcc phages which had positive *GRC*_*x*_, implying that they have at least some antibacterial activity, we found that overall, the 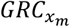 increases with the MOI (**Supplementary Figure 3**). A similar relationship was identified mathematically between MOI and VI for *Escherichia coli* phages T4, T5 and T7 under all tested conditions, as well as between MOI and qualitatively assessed growth curve reduction for *Salmonella enterica* phage *ϕ*St1 and *Acinetobacter baumannii* phage KARL-1, possibly suggesting a widespread direct relationship between phage MOI and antibacterial activity (25, 57, 58). Since this finding suggests that Bcc phages could be useful at high MOIs, even if their overall effectiveness across all MOIs is low, we considered both the *GRC*_*x*_ and the 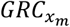 at the maximum available MOI – designated 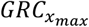 – when evaluating the potential therapeutic usefulness of Bcc phages. Seeking to establish a guideline with which to determine if Bcc phages have therapeutic potential, we set a GRC threshold of *GRC*_*x*_ *<u>or</u>* 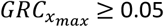, meaning that a phage must reduce bacterial growth by at least 5% – through either of these metrics – in order to be considered therapeutically useful. Values for *GRC*_*x*_ and 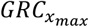 are summarized for all tested phage-host pairs in **Figure 2**, demonstrating substantial variability for both metrics among Bcc-targeting phages.

**Figure 1:**
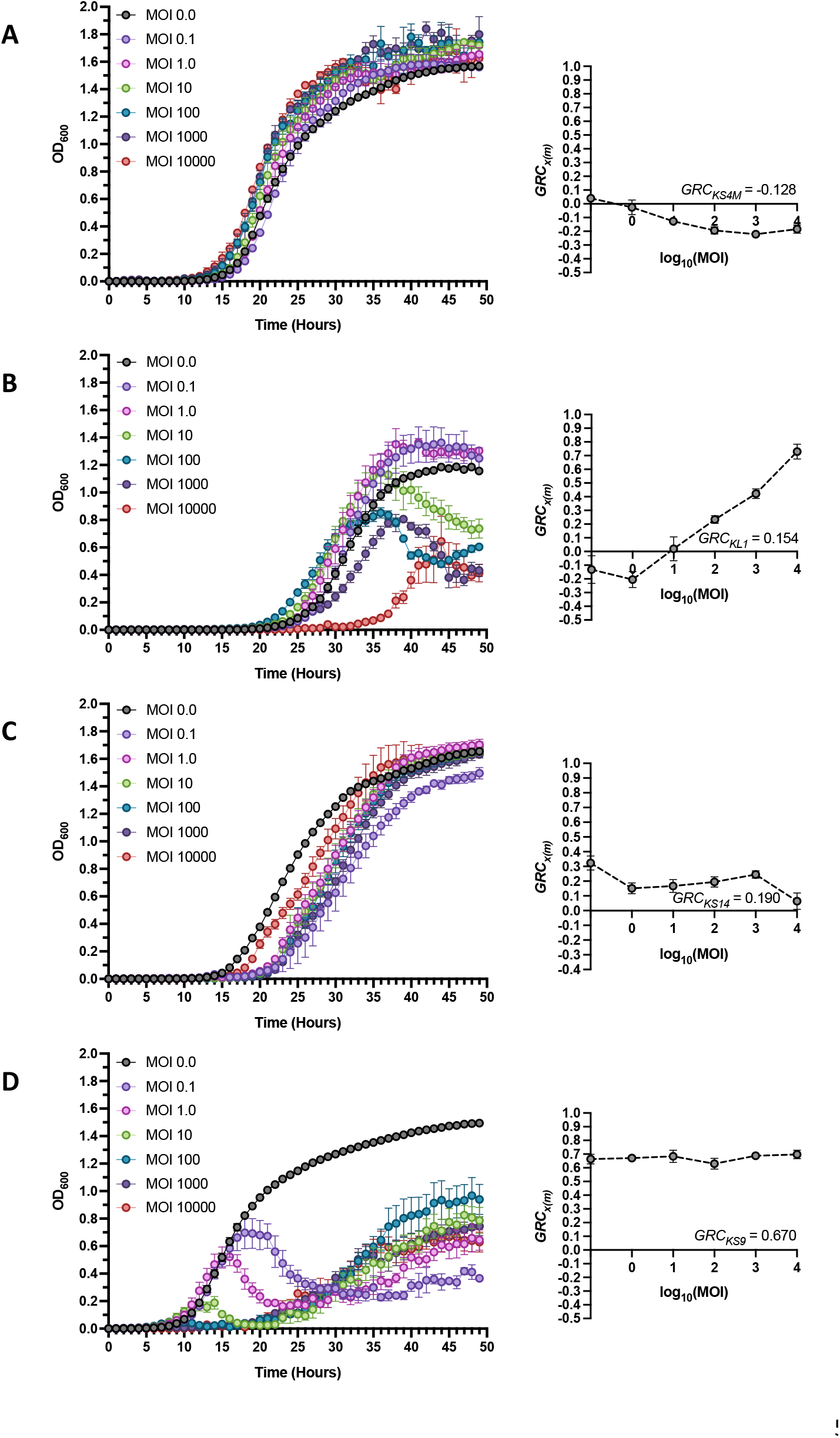
Representative Growth Reduction Trends of Phages Targeting the Bcc. Bacterial growth curves (left) and associated Growth Reduction Coefficient (GRC) curves (right) of various Bcc phage-host pairs for infections conducted at standard conditions across the standard MOI range of 0.1 to 10000. Displayed phage-host pairs are representative of situations in which the established GRC threshold *GRC*_*x*_ or 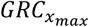 ≥ 0.05 is not met (**A**; KS4M on B. cenocepacia C6433), and of MOI-GRC trends in which the 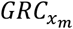 increases with MOI (**B**; KL1 on B. cenocepacia AU41264), in which the 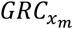 decreases with MOI (**C**; KS14 on B. multivorans C5393), and in which the 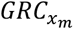 appears to be relatively independent of the MOI (**D**; KS9 on B. cenocepacia AU41386). Black lines represent bacterial growth without phage (MOI 0.0), while coloured lines represent each of the investigated MOIs. 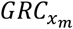 values were computed for each MOI using **equation 3**, and the *GRC*_*x*_ values for all representative phage-host pairs were then calculated using **equation 6** and are presented here for each representative pair. Bars represent standard error of the mean of at least three biological replicates.

**Figure 2:**
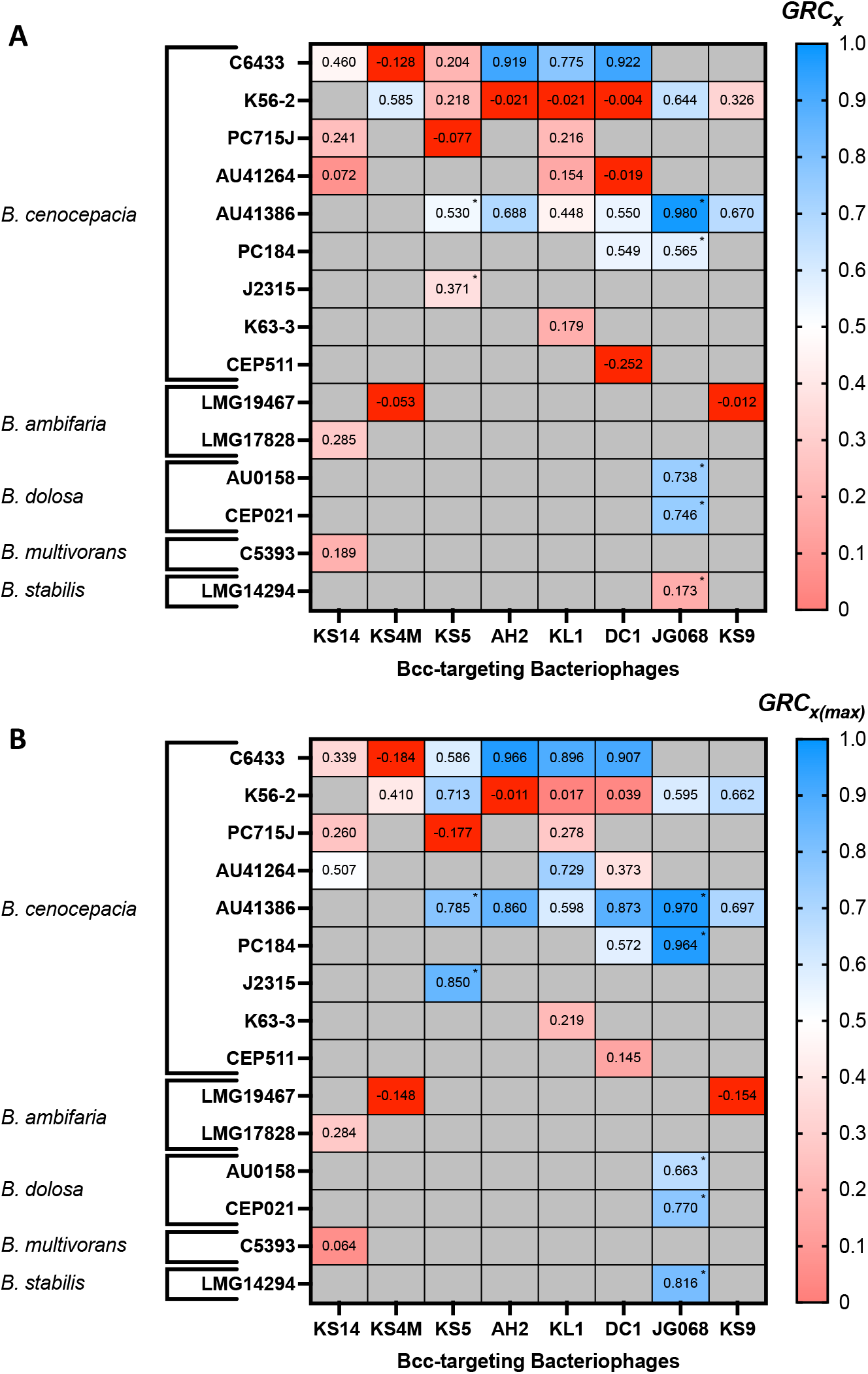
Growth Reduction Coefficients (GRC) of all available Bcc-phage host pairs. All infections were conducted at standard conditions across the standard MOI range of 0.1 to 10000, except for cells designated with * which indicate phage-host pairs where the maximum available MOI was 1000, and *GRC*_*x*_ (**A**) and 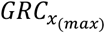 (**B**) values were then computed for each phage-host pair using **equations 6** and **3**, respectively. Data are presented at heatmaps with high positive GRC values in blue and low positive GRC values in pink, while negative values are shown in red, and GRC values are also provided numerically in each cell. Cells shaded in gray represent phage-host pairs which did not satisfy the EPA threshold and were not investigated. Values represent the mean of at least three biological replicates.

Among the 37 phage-host pairs investigated, 7 failed to pass the GRC threshold since neither their *GRC*_*x*_ or 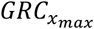 values reached 0.05 (representative example: phage KS4M on *B. cenocepacia* strain C6433; **Figure 1A**). Interestingly, all seven of these pairs have negative *GRC*_*x*_, suggesting that these phages increase bacterial growth rather than reducing it – implicating them as therapeutically unsuitable at these conditions. This phenomenon was not seen under any conditions for the phages investigated using the VI (25), implying that these are the first examples of phages being mathematically demonstrated to improve bacterial growth. Among the 30 remaining pairs, three distinct 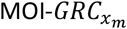 trends were identified. A direct relationship between these variables, as described previously and in other studies (25, 57, 58), was seen in 16 of these pairs (representative example: phage KL1 on *B. cenocepacia* strain AU41264; **Figure 1B**), while a modest inverse relationship was observed for only 2 pairs (representative example: phage KS14 on *B. multivorans* strain C5393; **Figure 1C**). Interestingly, in 6 of the pairs the 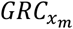 of the phage did not change significantly across the tested MOI range, suggesting that in these cases the GRC is independent of the MOI (representative example: KS9 on *B. cenocepacia* strain AU41386; **Figure 1D**). In the remaining 6 pairs, the relationships between MOI and 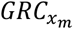 are irregular, meaning they fit into none of the patterns identified above. In some of these cases, such as phage KL1 on *B. cenocepacia* strain C6433 (see **Supplementary Figure 2C**), the overall 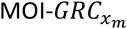 trend is positive but unusual features – such as vertices at intermediate MOIs – prevent their classification into the relevant category. Phages exhibit highly distinct 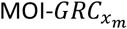 trends (see **Supplementary Figure 3**), in addition to highly distinct 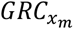 values, on different bacterial strains – indicating that the specific phage-host pair is crucial to whether a phage might be therapeutically effective. These highly variable findings highlight the enormous diversity in the antimicrobial effects of both LC and OL Bcc phages, and underscore the need to evaluate these phages against particular hosts on a case-by-case basis. Moreover, we found that efficiency of phage activity (EPA), is a poor predictor of the antibacterial effects of phages against planktonic cells, as measured by GRC (**Supplementary Figure 4**). Specifically, although modest correlations were observed between EPA and both *GRC*_*x*_ and 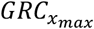 (see **Supplementary Table 3**), and a low EPA is reasonably predictive of low *GRC*_*x*_ and 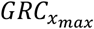, a high EPA is by no means predictive of high *GRC*_*x*_ or 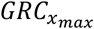. As a result, although low EPA may be used to disqualify certain phage-host pairs, a high EPA cannot be used to suggest that the phage might be therapeutically effective against that specific host.

### The Endpoint Growth Reduction Coefficient inaccurately approximates the Growth Reduction Coefficient

Although approaches such as the VI or GRC, which use data for growth over an extended period of time, are obviously more informative than using growth data from the single endpoint of that growth period, the former techniques are either costly or time consuming depending on whether or not modern instruments are available, and using an endpoint measurement as an approximation may therefore be tempting (58, 59). To investigate whether such an approximation might be valid, we defined an endpoint Growth Reduction Coefficient (eGRC) which is similar to the GRC but uses a single endpoint growth measurement rather than the area under a continuous growth curve. In this approach, a target bacterial strain is grown in the presence and absence of a phage *x* at various multiplicities of infection (MOI, *m*), and we utilize the difference between the final and initial growth values to define the endpoint Growth, 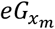, as shown in **equation 7**:

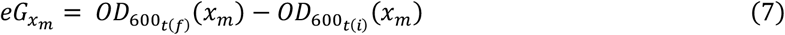

The endpoint Growth Reduction Coefficient of phage *x* at 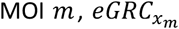, is then defined through comparison with the endpoint Growth of the untreated bacterial strain, *eG*_0_, as shown in **equation 8**:

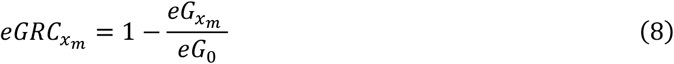

Using this approach, 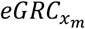 values were computed at all available MOIs for all Bcc phage-host pairs for which an EPA ≥ 10^−5^ was obtained, and were plotted against their cognate 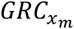 values to determine the similarity between the two values (**Figure 3**). We found that the absolute difference between the 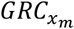 and 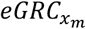 values was 0.10 or less (≤10% of GRC metric) in only 40.93% of cases, meaning there is a substantial difference between these two values in a majority of cases. Among plotted points which had a 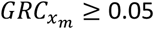, meaning they can be considered therapeutically relevant according to our criteria, the absolute difference between 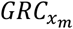 and 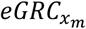 was 0.10 or less in only 30.26% of cases. In 13.82% of cases the 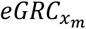 was substantially larger than the 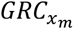, while in the remaining 55.92% of cases the 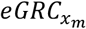 was substantially smaller than the 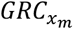. These findings suggest that in general, endpoint measurements such as 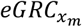 provide inaccurate approximations of growth-curve based approaches like 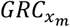 because they severely underestimate the true growth reduction capabilities of phages, and should therefore be used only sparingly in situations where growth-curve based approaches are impossible.

**Figure 3:**
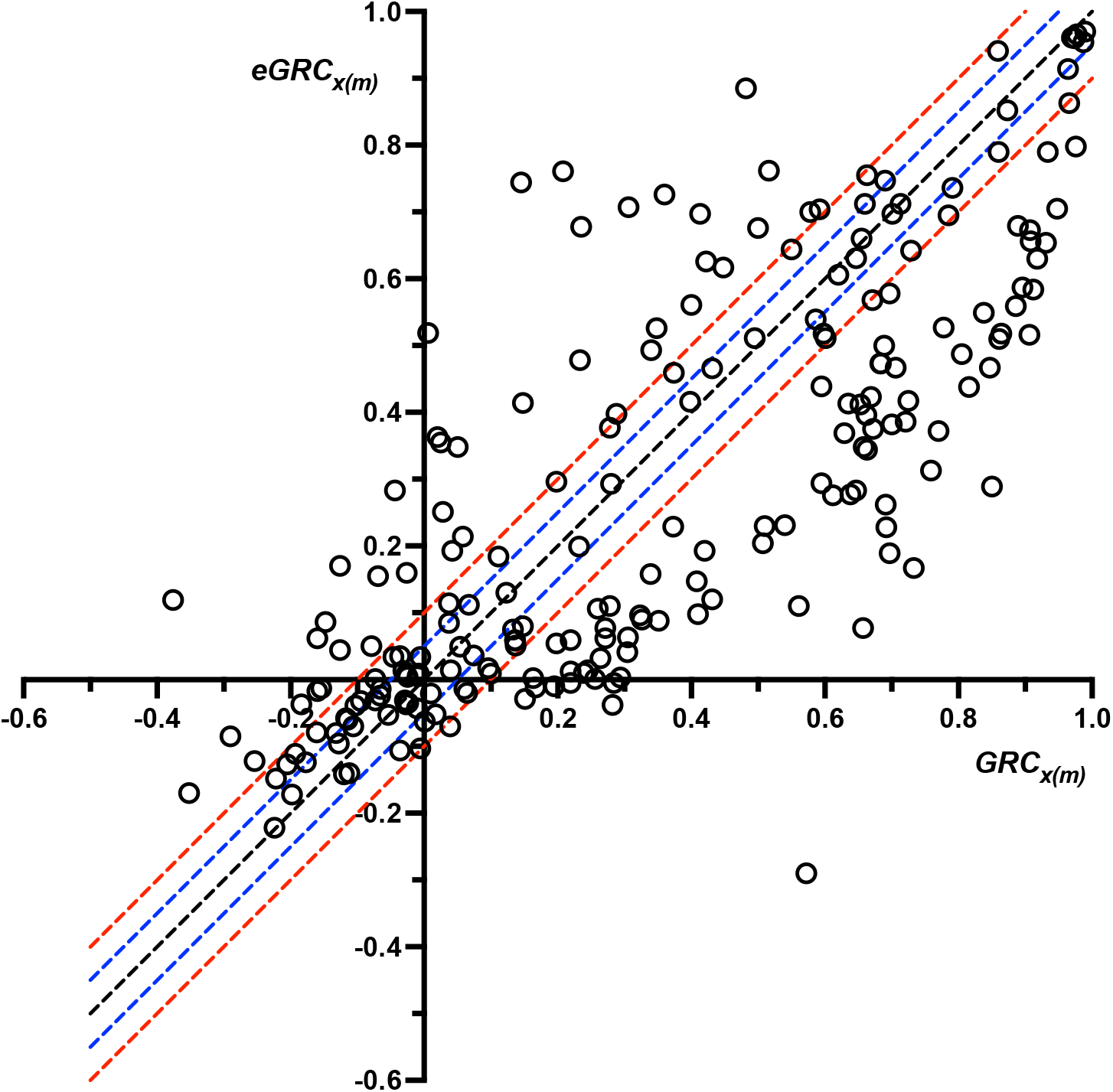
Comparison of 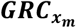 and 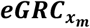 of Phages Targeting the Bcc. 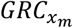 and 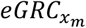 values of all investigated Bcc phage-host pairs, for infections conducted at standard conditions across the standard MOI range of 0.1 to 10000, were calculated using **equations 3** and **8**, respectively, and were plotted using an x-y scatterplot. The absolute difference between the two values was computed for each phage-host pair, and was used to determine the accuracy with which 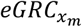 can estimate the 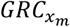. If 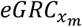 were a perfect predictor of 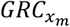, all plotted points should theoretically fall on the black dashed mirror line, which represents identical 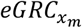 and 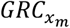 values. Points which fall above the mirror line are cases where the 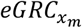 overestimates the 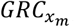, while those which fall below the mirror line are cases where the 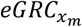 underestimates the 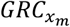. The blue and red dashed lines denote absolute 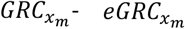 differences of 0.05 (5% of GRC) and 0.10 (10% of GRC), respectively, meaning that points which occupy the region between the blue lines, inclusive, have an absolute 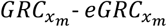 difference of 0.05 or less while points which occupy the region between the red lines, inclusive, have an absolute 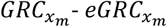 difference of 0.10 or less. Plotted points represent the mean values of at least three biological replicates.

### Lysogenization Frequency (*f*_(*lys*)_), a novel metric for quantification of a phage’s tendency to form stable lysogens, reveals enormous host and environment-driven diversity in the stable lysogen formation behaviours of Bcc phages

When considering the lysogenization behaviour of an LC phage in the context of its therapeutic suitability, the tendency of the phage to form unstable, short-lived lysogens is irrelevant since these lysogens will quickly be destroyed upon the phage’s reversion to the lytic cycle. Therefore, the important factor is the frequency with which the phage forms stable, long-term lysogens which contribute to bacterial growth and pathogenicity, which we call the lysogenization frequency (*f*_(*lys*)_). Concordantly, we define (*f*_(*lys*)_) as the proportion of survivors of infection by phage *x* (*s*_*x*_) which are lysogens of phage *x* (*l*_*x*_) after an extended period of time (*t*), at a particular set of environmental conditions including MOI (*m*), infection medium (*M*), temperature (*T*), and the host bacterium (*h*), as shown in **equation 9**:

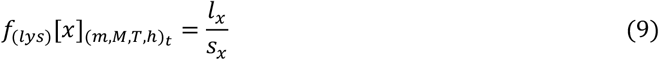

Since it can be demonstrated using the Poisson Distribution that every bacterium is infected by at least one phage in an infection at any MOI ≥ 8 (59), every bacterium recovered from an infection at such an MOI is by definition a survivor of infection, and the *f*_(*lys*)_ can therefore be estimated by screening these survivors for lysogeny. To investigate the effects of the infection medium, temperature, and the host organism on the *f*_(*lys*)_ of an LC phage, we quantified the *f*_(*lys*)_ of the *myovirus* KS14 in 48-hour liquid infections at the maximum available MOI (*f*_(*lys*)_ [*KS*14]_(*max*)_), on three strains of *B. cenocepacia* and one strain of *B. multivorans*, in several different infection media at various temperatures (**Figure 4A**). Interestingly, *f*_(*lys*)_ is consistently high in ½ LB medium in most of the tested temperature and host combinations, but varies substantially with both host and temperature on other media. Simultaneously, M9 MM is associated with lower *f*_(*lys*)_, possibly suggesting that lysogeny is favoured at higher nutrient densities among Bcc phages. Although the individual and combined effects of these environmental variables on *f*_(*lys*)_ are undoubtedly complex and a mechanistic understanding thereof remains elusive, particularly for the therapeutically relevant ACFSM and *Galleria mellonella* haemolymph, the observed diversity in *f*_(*lys*)_, even for a single phage, clearly demonstrates that the tendency of a phage to form stable lysogens is not determined solely by its genetic ability to do so. The *f*_(*lys*)_ should therefore be determined under relevant environmental conditions for any LC phage as part of its characterization and the determination of its suitability for therapeutic applications. Concordantly, we estimated the *f*_(*lys*)_ of all 8 Bcc phages on all available hosts, under the environmental conditions used in our GRC experiments, at the maximum available MOI (*f*_(*lys*)_ [*x*]_(*max*)_). Considerable variability in *f*_(*lys*)_ was observed between different Bcc phages as well as within individual phages, depending on the bacterial host (**Figure 4B**), further highlighting the importance of the specific phage-host pair to the phage’s tendency to form stable lysogens. Interestingly, many phage-host pairs produced no evidence of stable lysogeny despite the fact that all of these phages (excluding JG068) are genetically capable of forming lysogens. In some cases, this may be due to phage-host genetic incompatibility which prevents the integration of the phage genome into that of the host, while some strains may have other properties which prevent the establishment of lysogeny or cause rapid re-induction of the prophage to the lytic cycle. While the precise reasons behind these findings can only be speculated at present, it is clear that the stable lysogen forming behaviours of these genetically LC phages depend greatly on the particular strain being targeted, implying that at least some LC phages could potentially be used therapeutically against specific strains without treatment failure driven by the formation of lysogens.

**Figure 4:**
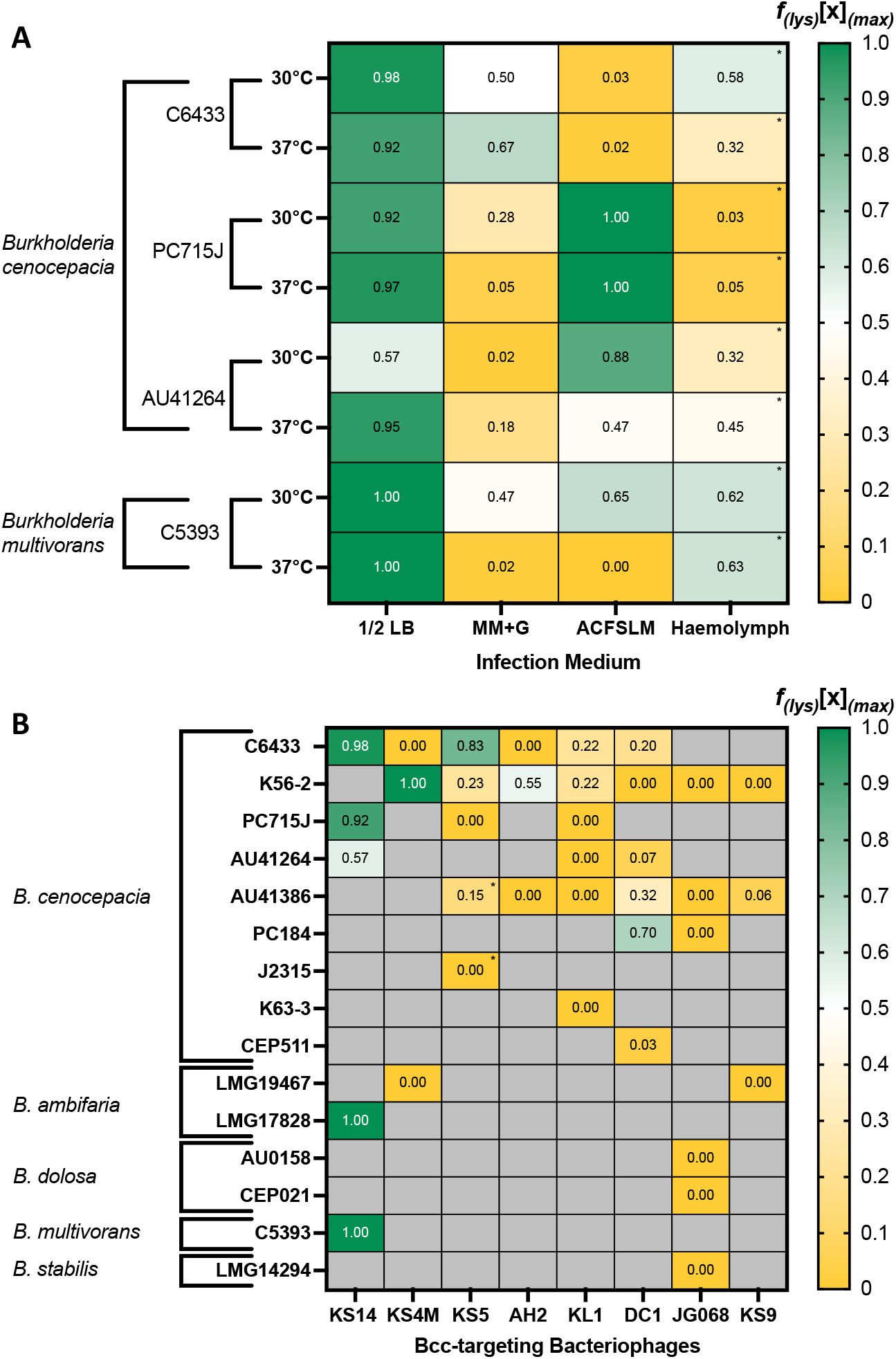
Lysogenization Frequencies (*f*_(*lys*)_) of Bcc-targeting phages. Infections were conducted in different experimental media and *in vivo* in the haemolymph of *Galleria mellonella* larvae, at different temperatures, and on different hosts with the Bcc-targeting myovirus KS14 (**A**), and at standard conditions for all Bcc phage-host pairs (**B**). All infections were performed at the maximum available MOI of 10000, except for cells designated with * which indicate phage-host pairs where the maximum available MOI was 1000, and *f*_(*lys*)_ values were then computed using **equation 9**. *f*_(*lys*)_ values for phage JG068 in (**B**) were assigned automatically as JG068 is obligately lytic and is genetically incapable of forming lysogens. Data are presented as heatmaps with high *f*_(*lys*)_ values in green and low *f*_(*lys*)_ in yellow, and *f*_(*lys*)_ values are also provided numerically in each cell. Cells shaded in gray represent phage-host pairs which did not satisfy the EPA threshold and were not investigated. Values represent the mean of at least four biological replicates with fifteen colonies tested for each biological replicate.

### Low-*f*_(*lys*)_ LC Bcc phages can be highly effective at reducing bacterial growth

Since many LC Bcc phages have low *f*_(*lys*)_ and may therefore engage in replication behaviour similar to that of OL phages, we sought to investigate whether these phages also have high growth reduction capabilities and might therefore be therapeutically useful. Concordantly, we compared the 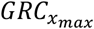 (**Figure 2B**) and *f*_(*lys*)_ [*x*]_(*max*)_ (**Figure 4B**) of all tested Bcc phage-host pairs (*n*=37) and found a strong inverse correlation between these variables among phage-host pairs for which at least modest antibacterial activity was observed (**Figure 5A**). A similar comparison between *f*_(*lys*)_ [*x*]_(*max*)_ (**Figure 4B**) and *GRC*_*x*_ (**Figure 2A**) found no significant correlation (**Supplementary Figure 5**; see **Supplementary Table 3** for details), which is unsurprising since *f*_(*lys*)_ likely varies with MOI. As previously described, 7 of these phage-host pairs failed to meet the established GRC ≥ 0.05 criterion, and no significant relationship between *f*_(*lys*)_ [*x*]_(*max*)_ and 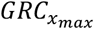 was identified for these pairs (**Figure 5A**, see **Supplementary Table 3** for details). Of the remaining 30 phage-host pairs, 26 (87%) followed a strong overall trend of *f*_(*lys*)_ [*x*]_(*max*)_ and 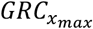 being inversely correlated (R^2^=0.67; *p<0*.*0001*), while another 4 pairs (13%) simultaneously had low *f*_(*lys*)_ [*x*]_(*max*)_ and 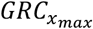 and thus form a small outlier group. Even when this outlier group is included, the inverse correlation between *f*_(*lys*)_ [*x*]_(*max*)_ and 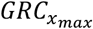 remains strong (R^2^=0.27; *p=0*.*0033*). Indeed, among the 30 phage-host pairs which satisfy the GRC ≥ 0.05 criterion, all of those with an *f*_(*lys*)_ [*x*]_(*max*)_ > 0.4 have a 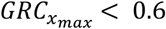, while 83% of those with a *f*_(*lys*)_ [*x*]_(*max*)_ < 0.4 have a 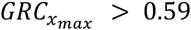, meaning that *f*_(*lys*)_ [*x*]_(*max*)_ is a fairly powerful predictor of 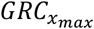 among these phages (**Figure 5A**). The presence of the outlier group suggests that a low (or absent) tendency to form stable lysogens does not invariably imply that a phage is effective at reducing growth – a finding which is further supported by the fact that 6 of the 7 phage-host pairs which fail to meet the GRC ≥ 0.05 criterion also have low *f*_(*lys*)_ [*x*]_(*max*)_. Importantly, this reiterates the fact that lysogeny is not the only obstacle for finding phages with strong antibacterial activity against certain host strains. Nevertheless, the fact that the vast majority of potentially therapeutically useful (GRC ≥ 0.05) Bcc phages follow a strong inverse correlation between *f*_(*lys*)_ [*x*]_(*max*)_ and 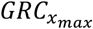 implies that high-*f*_(*lys*)_ [*x*]_(*max*)_ phages tend to have relatively low 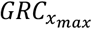 and are therefore unsuitable for monophage therapy, while low-*f*_(*lys*)_ [*x*]_(*max*)_ phages generally tend to have relatively high 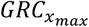 and could therefore be therapeutically useful. Crucially, several low-*f*_(*lys*)_ [*x*]_(*max*)_ LC phage-host pairs, such as those including LC phages AH2, KL1, or DC1, have 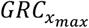 values comparable to (and in some cases higher than) those seen with the OL Bcc phage JG068 (**Figure 5A**, also see **Table 3**), strongly suggesting that low *f*_(*lys*)_ [*x*]_(*max*)_ LC phages such as these may be suitable for monophage therapy.

**Figure 5:**
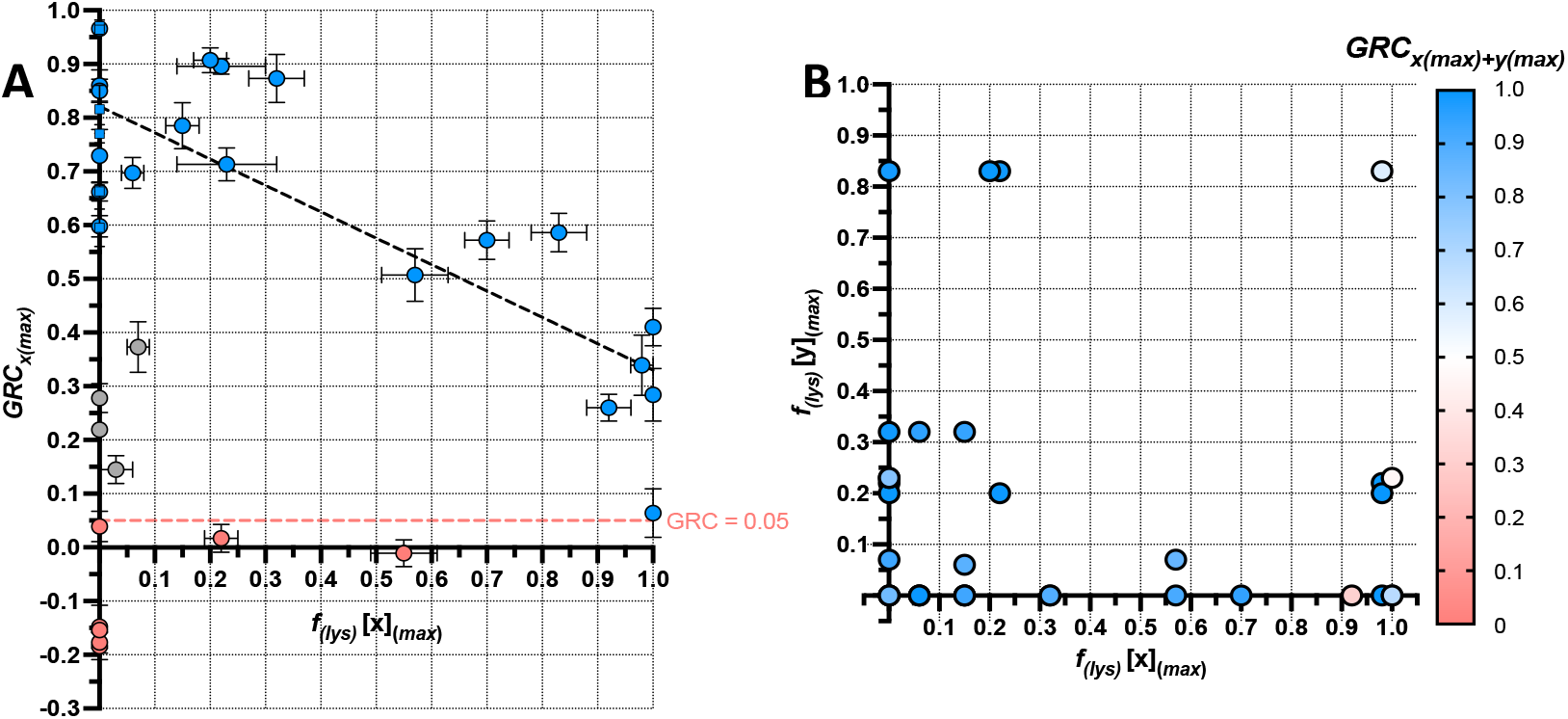
Growth Reduction Coefficient (GRC) as a function of Lysogenization Frequency (*f*_(*lys*)_) **(A)** Correlation between 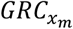 and *f*_(*lys*)_ at the maximum available MOI. 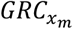 and *f*_(*lys*)_ values of all investigated Bcc single-phage-host pairs, for infections conducted at standard conditions at the maximum available MOI, were calculated using **equations 3** and **9**, respectively, and were plotted using an x-y scatterplot. Points shown in red represent phage-host pairs which did not satisfy the GRC ≥ 0.05 criterion, and thus fall below the red dashed line (representing a GRC of precisely 0.05). Point shown in grey represent phage-host pairs which did satisfy the GRC ≥ 0.05 criterion but form an outlier group which have low 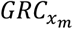 despite also having low *f*_(*lys*)_ values. Points shown in blue represent phage-host pairs which satisfied the GRC ≥ 0.05 criterion and follow the overall trend of 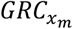 and *f*_(*lys*)_ values being inversely correlated. Circles and squares represent phage-host pairs containing LC and OL phages, respectively. The black dashed line represents the line-of-best-fit for points shown in blue. Vertical and horizontal error bars represent standard error of the mean of at least three and four biological replicates, respectively. **(B)** 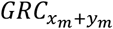 of two-phage cocktail-host pairs as a function of the *f*_(*lys*)_ values of the constituent phages of the cocktail, at the maximum available MOI. 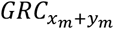 values of all possible 1:1 ratio two-phage cocktail-host pairs, along with the *f*_(*lys*)_ values of the constituent phages, were computed using **equations 3** and **9**, respectively, for infections conducted at standard conditions at the maximum available MOI. These values are plotted using a two dimensional scatterplot on which a third dimension, colour, is used to represent 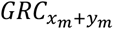. Plotted values represent the mean of at least three and four biological replicates for 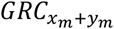 and *f*_(*lys*)_,respectively.

### High-*f*_(*lys*)_ LC Bcc phages can be highly effective at reducing bacterial growth when combined with low-*f*_(*lys*)_ LC or OL counterpart phages to form two-phage cocktails

Since high *f*_(*lys*)_ LC phages have low 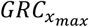 and are clearly unsuitable for therapeutic use on their own, we sought to investigate whether the therapeutic utility of such phages might be rehabilitated when they are combined with other phages. Concordantly, we constructed two-phage cocktails of all possible 1:1 ratio combinations of potentially therapeutically useful (GRC ≥ 0.05) Bcc phages and investigated their growth reduction effects, at maximum available MOI, on all *B. cenocepacia* strains for which more than one phage is available (**Figure 6**). Although the antibacterial effects of the two-phage cocktails, measured using 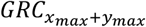, were in many cases higher than the antibacterial effects of either of the constituent phages on their own (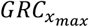 and 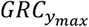), this effect was dependent on both the specific combination of phages being used and the host strain being targeted. Among cocktails of two low-*f*_(*lys*)_ LC phages and OL-low-*f*_(*lys*)_ LC phages, the magnitude of the improvement in antibacterial activity is highly variable, often because many of these phages are already highly effective in reducing bacterial growth on their own. The OL phage JG068, for instance, did not have substantially improved antibacterial activity when combined with any other phages targeting strains PC184 (**Figure 6E**) or AU41386 (**Figure 6C**) – likely because JG068 is already extremely effective against both of these strains on its own. Conversely, the antibacterial effect of JG068 is improved slightly when it is combined with any other phage targeting strain K56-2 (**Figure 6D**), on which its independent 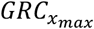 is lower. Similarly, the effectiveness of the low-*f*_(*lys*)_ LC phage AH2 is not significantly improved through combination with any phage on strains C6433 (**Figure 6A**) or AU41386 (**Figure 6B**) except phage KS9, which improves the antibacterial effect of AH2 on AU41386 substantially. Other low-*f*_(*lys*)_ LC phages such as DC1 and KL1 follow this trend as well, as they improve each other’s effectiveness on certain strains, such as C6433 (**Figure 6A**) and AU41264 (**Figure 6G**), but not on AU41386 (**Figure 6B**). Crucially, two-phage combinations containing a high *f*_(*lys*)_ LC phage paired with an OL or low-*f*_(*lys*)_ counterpart phage also have increased antibacterial effects relative to the effects of either of the constituent phages on their own. For instance KS14, which has high *f*_(*lys*)_ on all susceptible strains, improves the antibacterial effectiveness of all low-*f*_(*lys*)_ phages with which it is paired on strains C6433 (**Figure 6A**), PC715J (**Figure 6F**), and AU41264 (**Figure 6G**). Phage KS5 substantially improves the antibacterial activity of all low-*f*_(*lys*)_ phages on C6433 (**Figure 6A**), on which it has high *f*_(*lys*)_, but only modestly on K56-2 (**Figure 6D**), on which it has a relatively low *f*_(*lys*)_. However, KS5 also substantially improves the effectiveness of all low-*f*_(*lys*)_ phages on AU41386 (**Figure 6C**), on which it itself has low *f*_(*lys*)_, implying that the interactions between KS5 and other phages are, predictably, not affected solely by lysogenization Nevertheless, these results demonstrate that high-*f*_(*lys*)_ LC phages remain therapeutically useful since they can substantially improve the antibacterial effects of OL or low-*f*_(*lys*)_ phages with which they are combined. This finding is also evident when visualizing the antibacterial effect of the combination 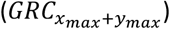 as a function of the *f*_(*lys*)_ of its constituent phages (**Figure 5B**). Although high-*f*_(*lys*)_ LC phages are unsuitable for monophage therapy since they produce only limited growth reduction when employed on their own (**Figure 5A**), cocktails in which such high-*f*_(*lys*)_ phages are combined with OL or low-*f*_(*lys*)_ LC counterpart phages consistently produce high levels of bacterial growth reduction (**Figure 5B**). Only two combinations of high-*f*_(*lys*)_-low-*f*_(*lys*)_ phages (KS4M & KS5 on K56-2, **Figure 6B**; KS14 & KL1 on PC715J; **Figure 6F**) produce poor antibacterial effects, and the low effectiveness of the constituent phages may be responsible in both cases. Only one cocktail of two high-*f*_(*lys*)_ phages (KS14 & KS5 on C6433, **Figure 6A**) was tested, and was found to have a limited ability to reduce bacterial growth – suggesting that when used in cocktails, high-*f*_(*lys*)_ phages should be paired only with low-*f*_(*lys*)_ or OL phages to achieve maximum antibacterial effect.

**Figure 6:**
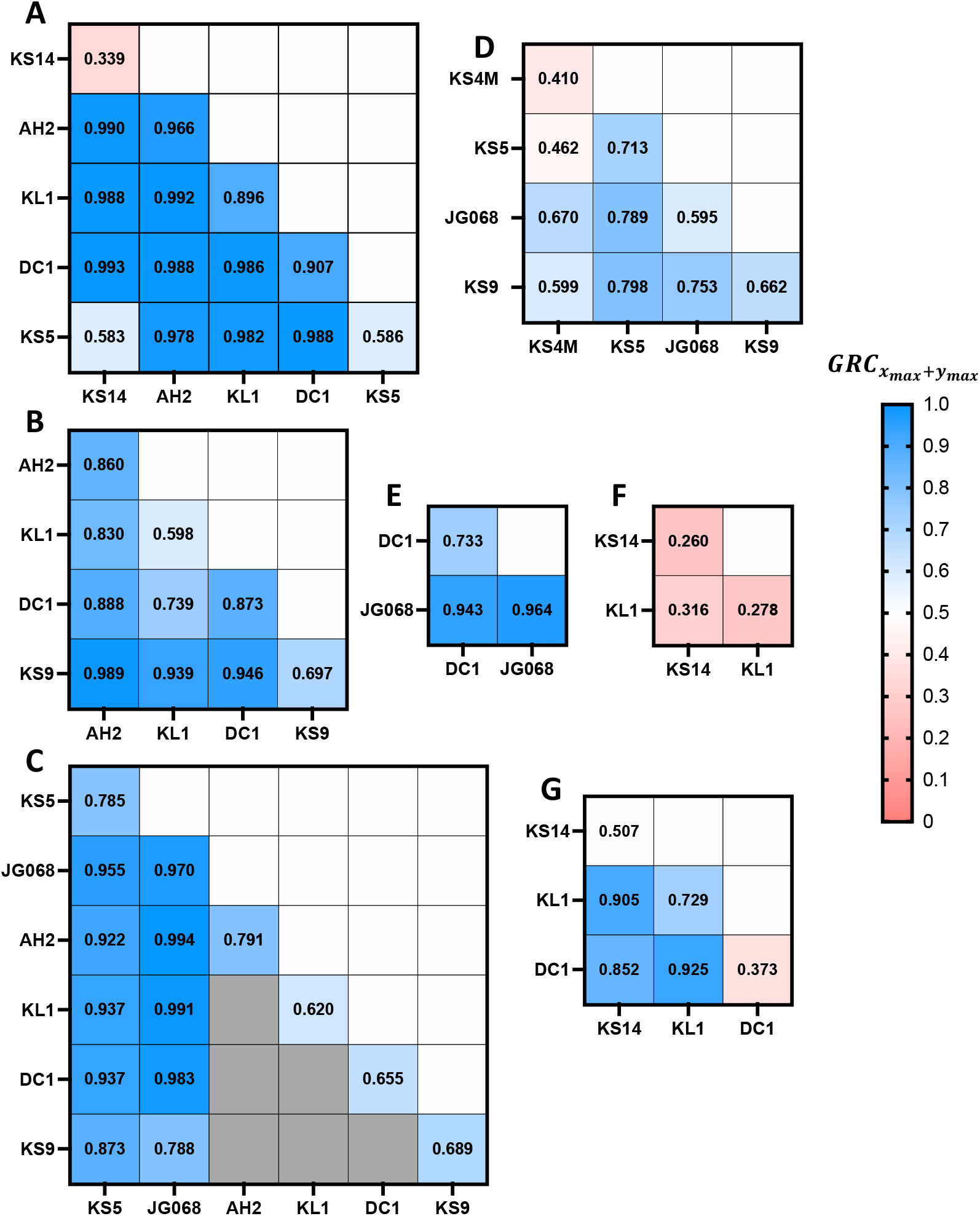
Growth Reduction Coefficients (GRC) at maximum available MOI for all available Bcc single-phage-host and two-phage cocktail–host pairs. 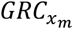 values for all Bcc single-phage-host pairs, as well as 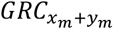 values for all possible 1:1 ratio two-phage cocktail-host pairs, for infections conducted at standard conditions at the maximum available MOI, were calculated using **equation 3**. Data are presented as heatmaps for all host strains for which more than one potentially therapeutically useful (GRC ≥ 0.05) phage is available, including *B. cenocepacia* strains C6433 (**A**), AU41386 at MOI: 10^4^ (**B**), AU41386 at MOI 10^3^ (**C**), K56-2 (**D**), PC184 (**E**), PC715J (**F**), and AU41264 (**G**). A gradient is used to colour code 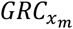 and 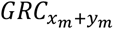 values within the heat maps, with high GRC values in blue and low GRC values in pink, and GRC values are also provided numerically in each box. Boxes shaded gray in (**C**) represent pairs which were not investigated at an MOI of 10^3^ because the maximum available MOI of those single-phage-host and thus two-phage cocktail-host pairs is 10^4^, and those pairs are displayed in (**B**). Values represent the mean of at least three biological replicates.

### LC Bcc phages interact synergistically to maximize reduction of bacterial growth

While many of the two-phage combinations described above appear to have antibacterial effects greater than the individual effects of each of their constituent phages (**Figure 6**), rigorous mathematical analysis is required to identify whether these differences are significant and whether they constitute synergistic interactions between phages or are merely the results of additive or independent effects. In studies investigating the effects of antibiotic combinations, independent or autonomous activity is defined, either qualitatively or quantitatively, as the antibacterial effect of combined treatment being indistinguishable from the effect of a single constituent of the combination – which suggests that this constituent is acting independently of the other agent and is responsible for the majority of the antimicrobial effect (61–64). Here we use the GRC metric to mathematically define autonomous activity in a cocktail of two phages *x, y* at a concentration *m* as a situation in which the GRC of the cocktail is statistically indistinguishable from the GRC of either of the constituent phages, as shown in **equation 10**:

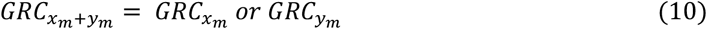

If an autonomous effect is not occurring, it can be surmised that both phages are contributing meaningfully to the antibacterial effect, and their effects must therefore be additive, synergistic, or antagonistic. If the antibacterial effects of the constituent phages are additive, we expect that the proportion of growth which remains after combined treatment should be statistically indistinguishable from the product of the proportions of growth which remain after treatment with each of the individual constituents (64–67). Using terms defined above, we express this mathematically as:

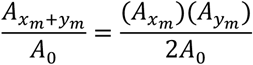

Rewriting this expression in terms of GRC, we define an additive effect as shown in **equation 11**:

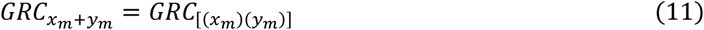

Where the right-side term 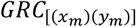 is used as a short form of 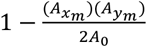. If the antibacterial interaction of the phages is synergistic, we expect that the growth reduction effect of the combination is significantly greater than the product of the individual antibacterial effects of the constituent phages, and we express this mathematically using GRC as shown in **equation 12**:

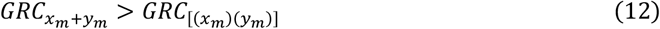

Conversely, it is possible that one of the phages has a much stronger antibacterial effect but the second phage reduces the effectiveness of the first phage. In such cases, the antibacterial interaction between the phages is antagonistic, and we expect the growth reduction effect of the combination to be less than the product of the individual antibacterial effects of the constituent phages (**equation 13**) and also less than the growth reduction effect of at least the more effective of the two phages (**equation 14**):

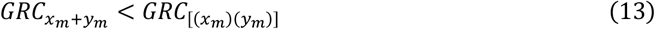

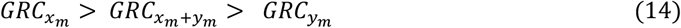

We conducted this mathematical analysis for all two-phage combinations which were tested against susceptible strains (*n*=36; see **Figure 6**), and identified six distinct types of antibacterial effects among these combinations (representative data in **Figure 7**, all data in **Supplementary Figure 6**, summarized in **Table 1**). Autonomous activity, defined in **equation 10**, was identified in 22 combinations (61.2%). In 18 (50.0%) of these, the overall antibacterial effect was driven by the more effective phage in the combination (representative example: JG068 & DC1 on PC184; **Figure 7A**). Many of these combinations involve the OL phage JG068 and the low-*f*_(*lys*)_ LC phage AH2 – which are extremely effective at reducing bacterial growth on their own and therefore there is little room for improvement through combination with another phage. Combinations tested against strain K56-2 (**Figure 6D**), which have modest and statistically insignificant improvements in antibacterial activity relative to their constituents, also fall into this category. In 2 (5.6%) combinations, the overall effect was driven by the less effective phage (**Supplementary Figure 6P** and **AD**), while the antibacterial effects of another 2 (5.6%) combinations were statistically indistinguishable from the effects of either of the constituents, likely because the constituents of these combinations were equally effective and ineffective at reducing growth (**Supplementary Figure 6E** and **AK**, respectively). Additive effects, defined in **equation 11** (representative example KS14 & KL1 on AU41264; **Figure 7B**), were identified for 7 (19.4%) combinations, all of these involving LC phages, and only 1 (2.7%) combination, KL1 and DC1 on AU41386 (**Figure 7C**), exhibited antagonistic effects as defined in **equations 13** and **14**. Synergistic interactions, defined in **equation 12** (representative example KS14 & DC1 on C6433; **Figure 7D**), were identified for 6 (16.7%) tested combinations. Crucially, all 6 of these synergistic combinations were composed of LC phages and, interestingly, 4 of these contained the high-*f*_(*lys*)_ LC phages KS14 or KS5. Although the strong antibacterial effects of these synergistically interacting phages suggest potential therapeutic utility, phage preparations for *in-vivo* use sometimes have lower MOIs than those used in these experiments, and we therefore sought to explore whether these synergistic interactions are preserved at lower MOIs. Concordantly, we tested our 6 synergistic phage combinations at MOIs of 10^0^ and 10^2^ (**Supplementary Figure 7**), and found that although powerful antibacterial effects are seen in the majority of these combinations even at lower MOIs, only four and two of the six combinations produced synergistic effects at MOIs 10^2^ and 10^0^, respectively, suggesting that these synergistic effects work best at higher MOIs. These findings are perhaps unsurprising considering that the majority of Bcc phages exhibit maximized antibacterial effects at higher MOIs (**Figure 1, Supplementary Figures 2 & 3**), and suggest that polyphage therapy of Bcc infections may require high MOIs for synergistic antibacterial effects. Interestingly, both DC1 and KS14 have positive effects on the growth of *B. cenocepacia* AU41264 at an MOI of 10^0^ and this effect translated to the combination of these phages as well, providing the only instance of a two-phage cocktail with negative 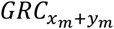 and thus a positive effect on bacterial growth (**Supplementary Figure 7I**). Nevertheless, synergistic antibacterial effects were preserved at lower MOIs for many phage combinations (**Supplementary Figure 7A-D,G,H**), possibly suggesting that MOI plays a role in the synergistic interactions of some phages but not others and thereby implying that different mechanisms of phage-phage synergy may be at play. Taken together, these findings demonstrate that both high and low-*f*_(*lys*)_ LC phages can greatly improve the antibacterial effectiveness of phage cocktails, often through mathematically proven synergistic interactions, and thus reveal a novel therapeutic role for LC phages.

**TABLE 1:** Synergistic, Additive, Antagonistic & Autonomous Growth Reduction Interactions between Bcc phages on susceptible strains of *Burkholderia cenocepacia*

| Host | Phage Pair | Conditions |  |  |  |  |  |  |  | Interpretation |
| --- | --- | --- | --- | --- | --- | --- | --- | --- | --- | --- |
| | | Autonomous Activity by Phage $x$? | | | Autonomous Activity by Phage $y$? | | | Interaction | | |
| | | $GRC(x_m + y_m)$ vs. $GRC(x_m)$ | | | $GRC(x_m + y_m)$ vs. $GRC(y_m)$ | | | $GRC(x_m + y_m)$ vs. $GRC[(x_m)(y_m)]$ | | |
|  |  | Outcome | Sign | Significance | Outcome | Sign | Significance | Sign | Significance |  |
| C6433 | $x$ = KS14<br>$y$ = KS5 | No | > | $p=0.0026$ | Yes | = | $p=0.9385(\text{ns})$ | < | $p=0.0007$ | Autonomous Activity |
| C6433 | $x$ = KS14<br>$y$ = AH2 | No | > | $p<0.0001$ | Yes | = | $p=0.0692(\text{ns})$ | = | $p=0.1739(\text{ns})$ | Autonomous Activity |
| C6433 | $x$ = KS14<br>$y$ = KL1 | No | > | $p<0.0001$ | No | > | $p<0.0001$ | > | $p=0.0008$ | Synergistic Interaction |
| C6433 | $x$ = KS14<br>$y$ = DC1 | No | > | $p<0.0001$ | No | > | $p=0.0057$ | > | $p=0.0097$ | Synergistic Interaction |
| C6433 | $x$ = KS5<br>$y$ = AH2 | No | > | $p<0.0001$ | Yes | = | $p=0.3591(\text{ns})$ | = | $p=0.2427(\text{ns})$ | Autonomous Activity |
| C6433 | $x$ = KS5<br>$y$ = KL1 | No | > | $p<0.0001$ | No | > | $p=0.0009$ | = | $p=0.1307(\text{ns})$ | Additive Activity |
| C6433 | $x$ = KS5<br>$y$ = DC1 | No | > | $p<0.0001$ | No | > | $p=0.0078$ | > | $p=0.0276$ | Synergistic Interaction |
| C6433 | $x$ = AH2<br>$y$ = KL1 | Yes | = | $p=0.0515(\text{ns})$ | No | > | $p<0.0001$ | < | $p=0.0264$ | Autonomous Activity |
| C6433 | $x$ = AH2<br>$y$ = DC1 | Yes | = | $p=0.0923(\text{ns})$ | No | > | $p=0.0078$ | < | $p=0.0005$ | Autonomous Activity |
| C6433 | $x$ = KL1<br>$y$ = DC1 | No | > | $p<0.0001$ | No | > | $p=0.0089$ | = | $p=0.5920(\text{ns})$ | Additive Activity |
| K56-2 | $x$ = KS4M<br>$y$ = KS5 | Yes | = | $p=0.5130(\text{ns})$ | No | < | $p=0.0347$ | < | $p=0.0182$ | Autonomous Activity |
| K56-2 | $x$ = KS4M<br>$y$ =JG068 | No | > | $p=0.0008$ | Yes | = | $p=0.1321(\text{ns})$ | = | $p=0.0583(\text{ns})$ | Autonomous Activity |
| K56-2 | $x$ = KS4M<br>$y$ = KS9 | No | > | $p=0.0030$ | Yes | = | $p=0.0949(\text{ns})$ | < | $p=0.0025$ | Autonomous Activity |
| K56-2 | $x = \text{KS5}$<br>$y = \text{JG068}$ | Yes | = | $p=0.1744(\text{ns})$ | No | > | $p=0.0089$ | = | $p=0.1061(\text{ns})$ | Autonomous Activity |
| K56-2 | $x = \text{KS5}$<br>$y = \text{KS9}$ | Yes | = | $p=0.1368(\text{ns})$ | No | > | $p=0.0466$ | = | $p=0.0925(\text{ns})$ | Autonomous Activity |
| K56-2 | $x = \text{JG068}$<br>$y = \text{KS9}$ | No | > | $p=0.0138$ | Yes | = | $p=0.0876(\text{ns})$ | = | $p=0.0608(\text{ns})$ | Autonomous Activity |
| PC715J | $x = \text{KS14}$<br>$y = \text{KL1}$ | Yes | = | $p=0.0863(\text{ns})$ | Yes | = | $p=0.2417(\text{ns})$ | < | $p=0.0024$ | Autonomous Activity |
| AU41264 | $x = \text{KS14}$<br>$y = \text{KL1}$ | No | > | $p=0.0003$ | No | > | $p=0.0157$ | = | $p=0.2727(\text{ns})$ | Additive Activity |
| AU41264 | $x = \text{KS14}$<br>$y = \text{DC1}$ | No | > | $p=0.0006$ | No | > | $p<0.0001$ | > | $p=0.0088$ | Synergistic Interaction |
| AU41264 | $x = \text{KL1}$<br>$y = \text{DC1}$ | No | > | $p=0.0143$ | No | > | $p<0.0001$ | = | $p=0.0980(\text{ns})$ | Additive Activity |
| AU41386 | $x = \text{KS5}$<br>$y = \text{AH2}$ | No | > | $p=0.0131$ | No | > | $p=0.0035$ | = | $p=0.0774(\text{ns})$ | Additive Activity |
| AU41386 | $x = \text{KS5}$<br>$y = \text{KL1}$ | No | > | $p=0.0175$ | No | > | $p=0.0070$ | = | $p=0.6356(\text{ns})$ | Additive Activity |
| AU41386 | $x = \text{KS5}$<br>$y = \text{DC1}$ | No | > | $p=0.0104$ | No | > | $p=0.0002$ | = | $p=0.7014(\text{ns})$ | Additive Activity |
| AU41386 | $x = \text{KS5}$<br>$y = \text{JG068}$ | No | > | $p=0.0072$ | Yes | = | $p=0.6379(\text{ns})$ | = | $p=0.2716(\text{ns})$ | Autonomous Activity |
| AU41386 | $x = \text{KS5}$<br>$y = \text{KS9}$ | Yes | = | $p=0.1078(\text{ns})$ | No | > | $p=0.0040$ | = | $p=0.1206(\text{ns})$ | Autonomous Activity |
| AU41386 | $x = \text{AH2}$<br>$y = \text{KL1}$ | Yes | = | $p=0.3590(\text{ns})$ | No | > | $p<0.0001$ | < | $p<0.0001$ | Autonomous Activity |
| AU41386 | $x = \text{AH2}$<br>$y = \text{DC1}$ | Yes | = | $p=0.5442(\text{ns})$ | Yes | = | $p=0.7920(\text{ns})$ | = | $p=0.0926(\text{ns})$ | Autonomous Activity |
| AU41386 | $x = \text{AH2}$<br>$y = \text{JG068}$ | No | > | $p=0.0010$ | Yes | = | $p=0.1039(\text{ns})$ | = | $p>0.9999(\text{ns})$ | Autonomous Activity |
| AU41386 | $x = \text{AH2}$<br>$y = \text{KS9}$ | No | > | $p=0.0066$ | No | > | $p<0.0001$ | > | $p=0.0261$ | Synergistic Interaction |
| AU41386 | $x = \text{KL1}$<br>$y = \text{DC1}$ | No | > | $p=0.0001$ | No | < | $p=0.0179$ | < | $p<0.0001$ | Antagonistic Interaction |
| AU41386 | $x = \text{KL1}$<br>$y = \text{JG068}$ | No | > | $p=0.0054$ | Yes | = | $p=0.1542(\text{ns})$ | = | $p=0.7879$ | Autonomous Activity |
| AU41386 | $x = \text{KL1}$<br>$y = \text{KS9}$ | No | > | $p<0.0001$ | No | > | $p<0.0001$ | > | $p=0.0063$ | Synergistic Interaction |
| AU41386 | $x = \text{DC1}$<br>$y = \text{JG068}$ | No | > | $p<0.0001$ | Yes | = | $p=0.4741(\text{ns})$ | = | $p=0.6263(\text{ns})$ | Autonomous Activity |
| AU41386 | $x = \text{DC1}$<br>$y = \text{KS9}$ | Yes | = | $p=0.1452(\text{ns})$ | No | > | $p<0.0001$ | = | $p=0.3185(\text{ns})$ | Autonomous Activity |
| AU41386 | $x = \text{JG068}$<br>$y = \text{KS9}$ | No | < | $p=0.0207$ | Yes | = | $p=0.0728(\text{ns})$ | < | $p=0.0240$ | Autonomous Activity |
| PC184 | $x = \text{DC1}$<br>$y = \text{JG068}$ | No | > | $p=0.0033$ | Yes | = | $p=0.2289(\text{ns})$ | = | $p=0.0519(\text{ns})$ | Autonomous Activity |

**Figure 7:**
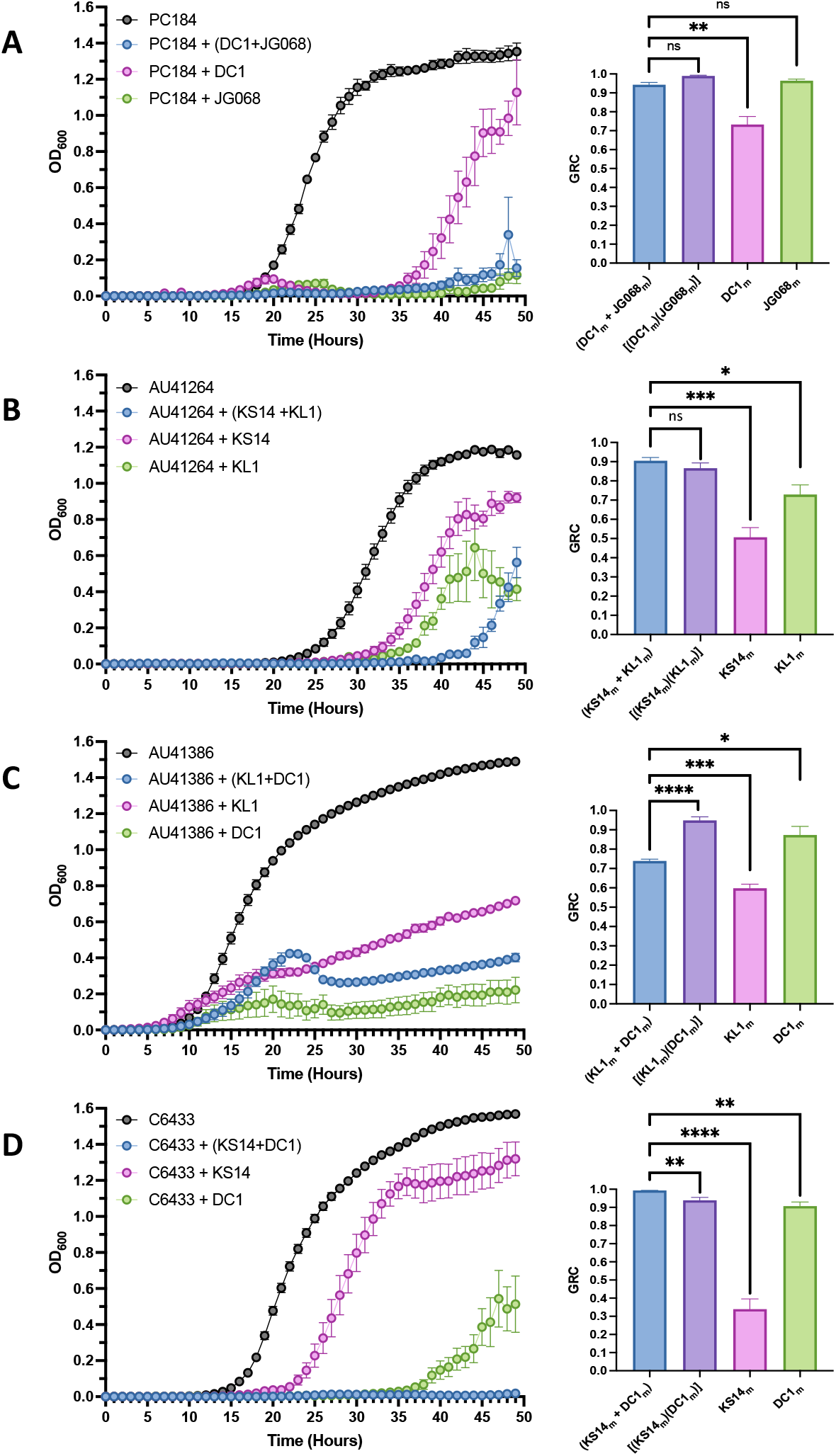
Representative Interactions between Bcc-targeting Phages. Bacterial growth curves (left) and treatment GRC bar charts (right) representative of two-phage combination-host pairs in which the phages behave **(A)** autonomously: where the effectiveness of the combinations is statistically indistinguishable from the effectiveness of one of the phages alone, **(B)** additively: where the effectiveness of the combination is statistically indistinguishable from the product of the individual effectivenesses of the component phages, **(C)** antagonistically: where the effectiveness of the combination is statistically lesser than the product of the individual effectivenesses of the component phages, or **(D)** synergistically: where the effectiveness of the combination is statistically larger than the product of the individual effectivenesses of the component phages. Purple bars represent the product of the individual effectivenesses of the component phages, as computed using the right-hand side of **equation 11**. Black line represents bacterial growth without phage. Blue lines and bars represent bacterial growth with and GRC of the phage pair, while pink and green represent bacterial growth with and GRC of the individual phages. Bars represent standard error of the mean of at least three biological replicates. **** indicates *p* < 0.0001, *** indicates *p* < 0.001, ** indicates *p* < 0.01, * indicates *p* < 0.05, and ns indicates *p* > 0.05.

## Discussion

Although phage therapy is a promising potential solution to the global threat posed by AMR, the current paradigm of exclusively using obligately lytic (OL) phages severely restricts the range of organisms against which phage therapy can be utilized, simply due to the fact that OL phages appear to be rare for many pathogenic species – including the members of the Bcc. In this study, we first address this issue from a theoretical perspective by proposing the more accurate term Lysogenization-Capable (LC) to describe phages which have the genetic capacity to form lysogens – thereby emphasizing the fact that the mere capability to form stable lysogens is not the sole predictor of whether or not a phage will do so under therapeutically-relevant conditions, which in turn implies that at least some LC phages may not form stable lysogens under such conditions and may therefore have therapeutic utility. We then proceed to examine this concept by introducing several novel metrics – Efficiency of Phage Activity (EPA), Growth Reduction Coefficient (GRC) and Lysogenization Frequency (*f*_(*lys*)_) – with which to quantify, in an *in-vitro* setting, the potential therapeutic suitability of both LC and OL phages.

EPA, a modified form of the canonical EOP metric, gauges phage activity on solid medium by taking into account both plaques and non-plaque evidence of phage activity – including mottling and turbid clearance, which can be due to low-level productive infection but also non-productive activity such as abortive infection (52) or lysis from without (53). Although these non-productive activities may be therapeutically suboptimal, using the EPA allows for a quantitative characterization of the effects certain phages have on particular hosts through these mechanisms, as well as through productive infections, which decreases reliance on qualitative and thus subjective approaches such as the direct spot test (51, 68). EPA therefore serves as a preliminary screen through which to eliminate phage-host pairs for which very low (or no) activity is recorded, and select more promising pairs which are then investigated using more sophisticated approaches such as the GRC. The GRC, a modified form of the recently devised Virulence Index (25), measures the ability of a phage to reduce bacterial growth in a planktonic killing assay (PKA) across a therapeutically relevant period of time, rather than only during the logarithmic phase – thereby capturing the ability of the phage (or cocktail) to prevent outgrowths which can occur at later timepoints. In this study, we utilized an EPA ≥ -5 threshold to exclude phage-host pairs with very low activity, but subsequently found that even pairs with EPA ≤ -4 are associated with low GRCs (**Supplementary Figure 4**), suggesting that a more restrictive threshold might be advisable in future studies. Importantly, high EPA is not a good predictor of high GRC, meaning that while the EPA screening approach can be used to eliminate phage-host pairs with low activity, it cannot be used to suggest that high EPA pairs necessarily show therapeutic potential (**Supplementary Figure 4**). Substantial diversity in EPA, GRC and MOI-GRC trends was noted among Bcc phages both between distinct phages and within individual phages depending on the specific host being targeted (**Figures 1 & 2**; **Supplementary Figures 1 & 2**). These differences might be the result of differences in infection parameters such as phage-host affinity, infectivity, burst size, and time to lysis (69), but could also be due to host factors such as receptor availability and phage resistance (8, 52). To further emphasize the importance of growth curve-based approaches such as the VI and GRC, we devised a derivative of the GRC termed endpoint GRC (eGRC) – which measures the growth reduction capacity of a phage using only the endpoint of bacterial growth – and found that this approach severely underestimates the GRC (**Figure 3**), meaning that endpoint approaches should only be used in circumstances where growth curve-based methods are not possible. *f*_(*lys*)_ is a novel variable which gauges the stable-lysogen formation behaviours of LC phages, and the observed variation in this metric is crucial to our proposed paradigm of phages occupying points on a spectrum in terms of their tendency to form stable lysogens. Indeed, the *f*_(*lys*)_ of LC phages appears to vary widely with environmental factors such as temperature, infection medium, and host, and varies substantially between different LC phages as well (**Figure 4**), which supports our conjecture that the genetic capacity to form lysogens is only a small part of the overall picture. Curiously, many LC phages fail to produce any evidence of stable lysogen formation on particular hosts, which may be due to genetic incompatibilities between the *attP* and *attB* sites of the phages and hosts, respectively. Other phages interestingly exhibit low (but non-zero) *f*_(*lys*)_ on certain hosts, which could be caused by “leaky” integration sites which lead to higher rates of spontaneous prophage induction, or perhaps by bacterial physiological responses which interfere with the establishment or maintenance of lysogeny in some strains (26, 27). The *f*_(*lys*)_ of the myovirus KS14 appears to be higher in nutrient-rich ½ LB and lower in M9 MM, possibly suggesting that lysogeny is favoured under high-nutrient conditions in Bcc phages – a trend which is opposite to that described for the coliphage lambda (26) – but the overall effects of temperature and infection medium on *f*_(*lys*)_ are challenging to interpret and require further investigation.

Although the mechanistic underpinnings of the enormous variability in EPA, GRC, and *f*_(*lys*)_ reported among Bcc phages remains, for the most part, difficult to understand, the fact that such variability exists, particularly among LC phages, reiterates the fact that specific environmental and host conditions are key determinants of whether a phage behaves in a therapeutically suitable manner. Concordantly, the potential therapeutic utility of LC phages must therefore be determined on a case-by-case basis through rigorous evaluation of their GRC and *f*_(*lys*)_, rather than through the overly simplistic, black-and-white approach of disqualifying phages based on the presence of a lysogenic cassette. In comparing the GRC and *f*_(*lys*)_ of Bcc phages, we found a highly significant (R^2^ = 0.67; *p*<0.0001) inverse relationship between these values for most phages which exhibit at least moderate (GRC ≥ 0.05) antibacterial activity (**Figure 5A**), demonstrating that low-*f*_(*lys*)_ LC Bcc phages can have high GRC values and may therefore be therapeutically suitable. Taken together, these findings illustrate the flawed nature of the current paradigm of exclusively using OL phages by revealing that LC phages vary substantially in their therapeutic suitability, and that while high-*f*_(*lys*)_ LC phages are unsuitable on their own, low*-f*_(*lys*)_ LC phages can be as effective as OL phages in monophage therapy.

Although monophage applications may be useful in situations where only one phage is available for a particular host, the high risk of resistance development – and the desire to avoid the strategic mistakes which led to the current AMR catastrophe – have led most researchers, in both experimental and clinical settings, to instead use treatments combining multiple phages simultaneously (8, 31, 70–74). Recognizing the therapeutic importance of these polyphage cocktails, we used the GRC and *f*_(*lys*)_ metrics to explore the therapeutic usefulness of LC Bcc phages in equal-ratio two-phage cocktails and, in particular, we sought to determine whether the effects of these two-phage cocktails result from significant, mathematically defined synergistic interactions. Although mathematically defined synergistic interactions have previously been reported for combinations of multiple antibiotics (61–63, 66) and combinations of single phages and antibiotics (64, 67), and several studies have demonstrated – without the use of rigorous mathematical definitions – improvements in antibacterial effect resulting from the combination of two or more phages (51, 75), to our knowledge this is the first study to report mathematically defined synergistic and antagonistic interactions, as well as autonomous and additive effects, between two phages. Interestingly, we found that many of these two-phage combinations produced powerful antibacterial effects which were either equal to (additive) or greater than (synergistic) the multiplicative antibacterial effects of their constituents (**Figure 6**; **Figure 7B & D**; all data summarized in **Table 1** and graphically in **Supplementary Figure 6**). Most importantly, neither low nor high-*f*_(*lys*)_ status in a constituent phage appears to be an impediment to the formation of cocktails with strong antibacterial activity, since all of the phages participating in additive or synergistic combinations were LC, and high-*f*_(*lys*)_ LC phages were in fact present in 4 of the 6 synergistic combinations identified (**Figure 5B**; **Table 1**) – including KS14 & DC1 on *B. cenocepacia* C6433, which had the strongest recorded antibacterial effect (**Figure 7D**). Furthermore, many of these combinations produce strong antibacterial effects even at lower MOIs (**Supplementary Figure 7**), meaning these synergistic effects could certainly be useful in therapeutic settings where lower MOIs are sometimes required. Taken together, these findings further suggest a shift from the current paradigm of exclusively using OL phages by revealing that LC phages may be rendered highly therapeutically useful through additive or synergistic interactions with other phages – which collectively produce powerful antibacterial effects. It is important to note, however, that this effect does not apply universally to all LC phages. On *B. cenocepacia* K56-2, for instance, KS4M (*f*_(*lys*)_= 1) fails to substantially improve the effectiveness of any phage with which it is paired, and in fact appears to reduce the antibacterial activity of KS5 (**Figure 6D**). Similarly, KS14 (*f*_(*lys*)_) = 0.92) does not improve the effectiveness of KL1 on PC715J (**Figure 6F**), suggesting that at least some LC phage pairing are therapeutically ineffective, and that a case-by-case evaluation of LC phage combinations on relevant hosts therefore remains indispensable.

When discussing the effects of polyphage cocktails, it is crucial to stress that although synergistic interactions are arguably the most interesting as they suggest interplays between individual phages, additive effects – so long as they substantially reduce bacterial growth – are equally useful from a therapeutic perspective. On the other hand, autonomous effects in which the more effective phage governs the overall antibacterial effect (**Figure 7A**) suggest that using the combination is not detrimental but provides no advantage relative to using the more effective phage on its own. Conversely, antagonistic interactions (**Figure 7C**) and autonomous effects in which the less effective phage governs the antibacterial interaction suggest that these combinations should be avoided, as they produce antibacterial effects which are lesser than those achievable through the use of the more effective phage alone.

Although a mathematical analysis of antibacterial effects is certainly crucial for determining whether the effects of a particular phage combination are autonomous, additive, synergistic, or antagonistic, it must be acknowledged that this approach does not facilitate a mechanistic understanding of how these effects occur. Although infection parameters such as phage-host affinity, infectivity, burst size and time to lysis may play a role in how phages can interact (25, 51, 69), these parameters remain largely unknown for Bcc phages. As well, whether the constituent phages of a cocktail utilize the same receptor could be useful for understanding their interactions, but primary receptors are known for only three of the eight Bcc phages examined in this study (**Supplementary Table 2**), rendering the interactions between these phages challenging to comprehend. For instance, in this study we report the first mathematically defined example of antagonism between two phages (**Figure 7C**), but the mechanism behind this interaction remains elusive. Similarly, the mechanisms whereby KS4M and KS9, despite being less effective than their counterpart phages, govern the antibacterial effects of their combinations with KS5 and JG068, respectively (**Supplementary Figure 6P & AD**), are also challenging to understand. Because KS4M has a high *f*_(*lys*)_, it is conceivable that superinfection immunity to KS5 by KS4M lysogens prevents the effectiveness of the former phage, possibly explaining the dominance of KS4M in this phage combination, but this explanation is unlikely for either of the other combinations described above since neither of those contain a high-*f*_(*lys*)_ phage. Autonomous effects where the more effective phage governs the overall effect, as well as additive effects and synergistic interactions, most likely result from combining phages which have particular infection parameters, as described previously, and the interplay between these parameters is likely most favourable in combinations of phages which interact synergistically (69). We suspect, however, that other factors may also be at least partially responsible for the synergistic interactions observed in this study. First, high-*f*_(*lys*)_ phages were present in four of these six combinations (see **Table 1**), raising the possibility that the formation of lysogens – which can be targeted by the other phage in the cocktail – could potentially play a role in the observed synergistic effect. Second, the other two combinations for which synergistic interactions were observed included phages which have distinct primary receptors (**Table 1**; **Supplementary Table 1**), raising the possibility of synergy due to decreased probability of simultaneous development of resistance to both of these phages (71). While tantalizing, these conjectures currently remain speculative and improved understanding of Bcc phage receptors, infection parameters, and the role of Bcc lysogens, is needed to shed light on the mechanisms behind these interactions.

In this study, we explored the effectiveness of two-phage combinations only at a ratio of 1:1, but our results suggest that GRC appears to be maximized at lower MOIs in a small minority of Bcc phage-host pairs (**Figure 1**; **Supplementary Figure 2**) - suggesting that combinations in which one phage is present in trace amounts relative to the other phage might produce ever stronger antibacterial effects in certain two-phage combinations. This idea was briefly explored by Storms et al., who found that 1:1 and 1:3 ratio combinations of the coliphages T7 and T5 had distinct VI values, although neither of these combinations was significantly different than monophage treatment with T7 (25). This suggests that future work in optimizing these polyphage combinations might focus on exploring the effects of ratios, by using the MOI-GRC trends identified in this study as a baseline for determining the MOIs at which the GRCs of individual Bcc phages are optimized.

Since the primary objective of this study was to assess the potential therapeutic suitability of LC Bcc phages, both in monophage and polyphage applications, based on *in-vitro* evaluation using our newly developed parameters, we did not focus on the effectiveness of these phages *in-vivo*. Although *in-vivo* polyphage treatment trials for Bcc infections have never been published, treatment of Bcc infections in *Galleria mellonella* larvae using combinations of antibiotics and single phages demonstrated antibacterial effects comparable to those seen in *in-vitro* phage-antibiotic anti-Bcc PKAs, possibly suggesting that polyphage treatments might have similar antibacterial effects *in-vivo* as they do *in-vitro* (32). Whether the synergistic interactions reported here might be maintained *in vivo* is difficult to predict, however, and warrants further investigation. Intriguingly, KS14 appears to be have a decreased *f*_(*lys*)_ in the haemolymph of *Galleria mellonella* (**Figure 4A**), meaning that if the synergistic interactions identified in several of our combinations are indeed at least partially reliant on the formation of lysogens – the magnitude of the synergistic effect may be diminished *in-vivo*. Conversely, synergistic interactions based on receptor identity would likely be preserved *in-vivo*, assuming that phage infectivity is unaffected under such conditions. Regardless, *in-vivo* polyphage treatments of Bcc infections are the next logical step in determining whether these phages may be used in human trials, and should be explored at length in subsequent studies.

Finally, efforts should be made to study the effects of phage-antibiotic cocktails containing high-*f*_(*lys*)_ LC phages and compounds which use activation of the cellular stress response to re-induce these phages to their lytic cycles. In a recent seminal study, Al-Anany et al. convincingly demonstrated that the LC coliphage HK97 can be therapeutically effective when paired with ciprofloxacin, since this compound triggers re-induction of the HK97 prophage to the lytic cycle, thereby improving overall bacterial clearance (33). Although we are not aware of any studies investigating combined treatments with LC phages and complement, this essential component of the innate immune system also stresses bacterial cells and could therefore improve overall growth reduction by triggering the re-induction of high-*f*_(*lys*)_ LC prophages. Crucially, the *f*_(*lys*)_ metric introduced in this study can be used to quantify the tendency of phages to form stable lysogens, and can thereby serve as an excellent screen for identifying phages which would likely synergize with prophage-inducing compounds.

In conclusion, it is reasonable to expect that the majority of all existing phages may be genetically capable of forming lysogens due to the evolutionary benefits of lysogenization – meaning that the existing paradigm of exclusively utilizing obligately lytic phages in phage therapy is therefore impractical. Viewed broadly, lysogenization among phages can pose a severe problem to the application of phage therapy, but this problem can be overcome if lysogenization-capable phages are used selectively and strategically. To that end, all LC phages must be evaluated for their tendency to form stable lysogens and their ability to reduce bacterial growth, using the *f*_(*lys*)_ and GRC metrics, respectively, as part of their initial characterization. Low-*f*_(*lys*)_ – High GRC phages may be used on their own if no alternatives are available, while both low and high-*f*_(*lys*)_ phages may be used in polyphage cocktails in which antibacterial activity is maximized through additive or synergistic interactions. These approaches can be increasingly elaborated by combining these phages with specific stress-inducing antibiotics, with the ultimate aim of constructing an LC polyphage-antibiotic cocktail which can completely abolish bacterial growth.

---

**Supplementary Figure 1:**
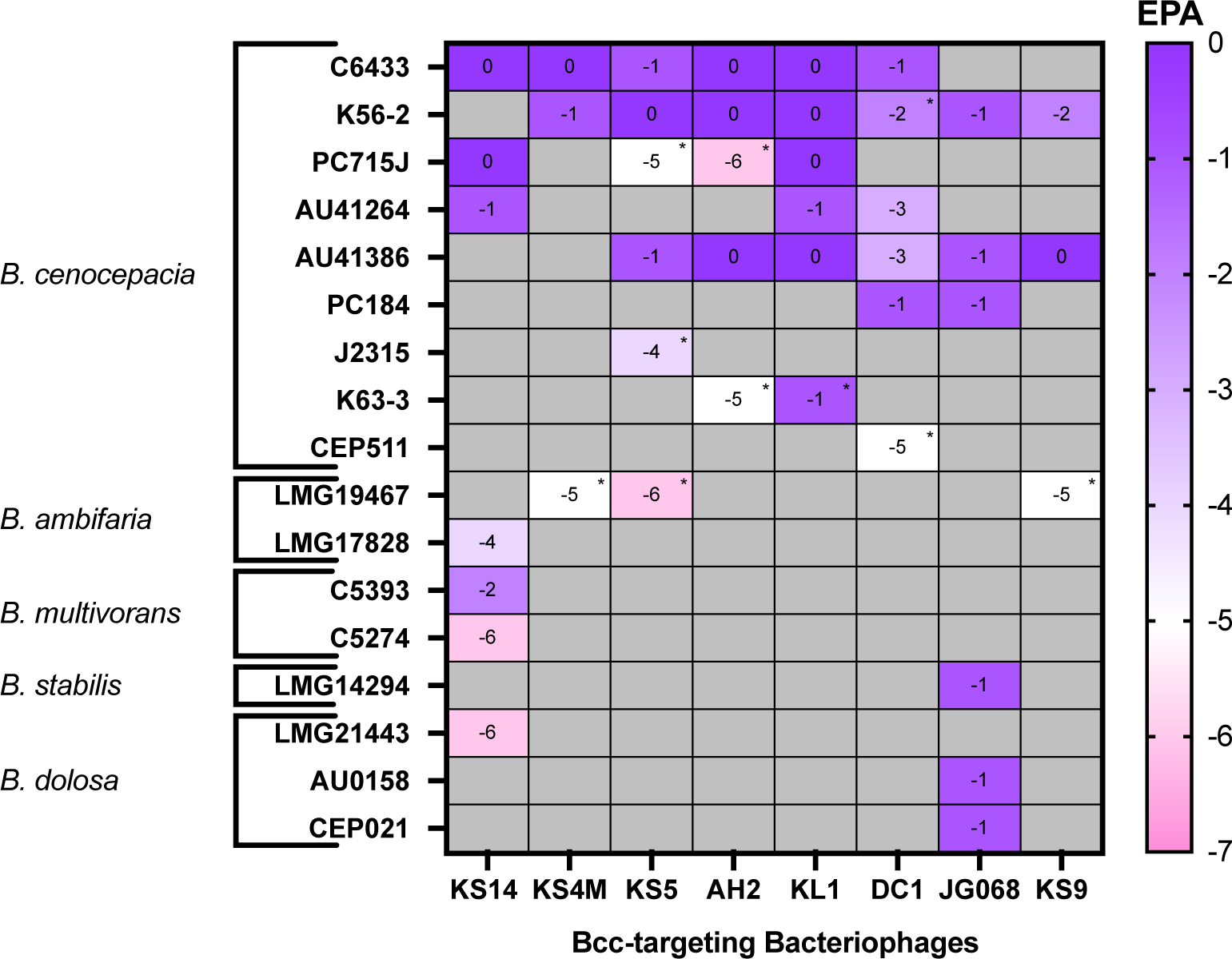
Efficiency of Phage Activity (EPA) of Bcc-targeting phages on susceptible Bcc species. Solid medium infections were conducted using a soft agar overlay serial dilution spotting assay, and EPA values were calculated for all phage-host pairs using **equation 1**. EPA values range from a maximum of 0 to a minimum of -7, while EPA < -7 indicates no phage sensitivity and is indicated by cells shaded in grey. Cells designated with * indicate phage-host pairs in which phages produce zones of bacterial lawn weakening or extremely turbid clearing, rather than discrete, readily countable plaques. An arbitrary threshold of EPA ≥ -5 was used to select phage-host pairs for further investigation, while phage-host pairs with EPA < -5 were excluded from further research.

**Supplementary Figure 2:**
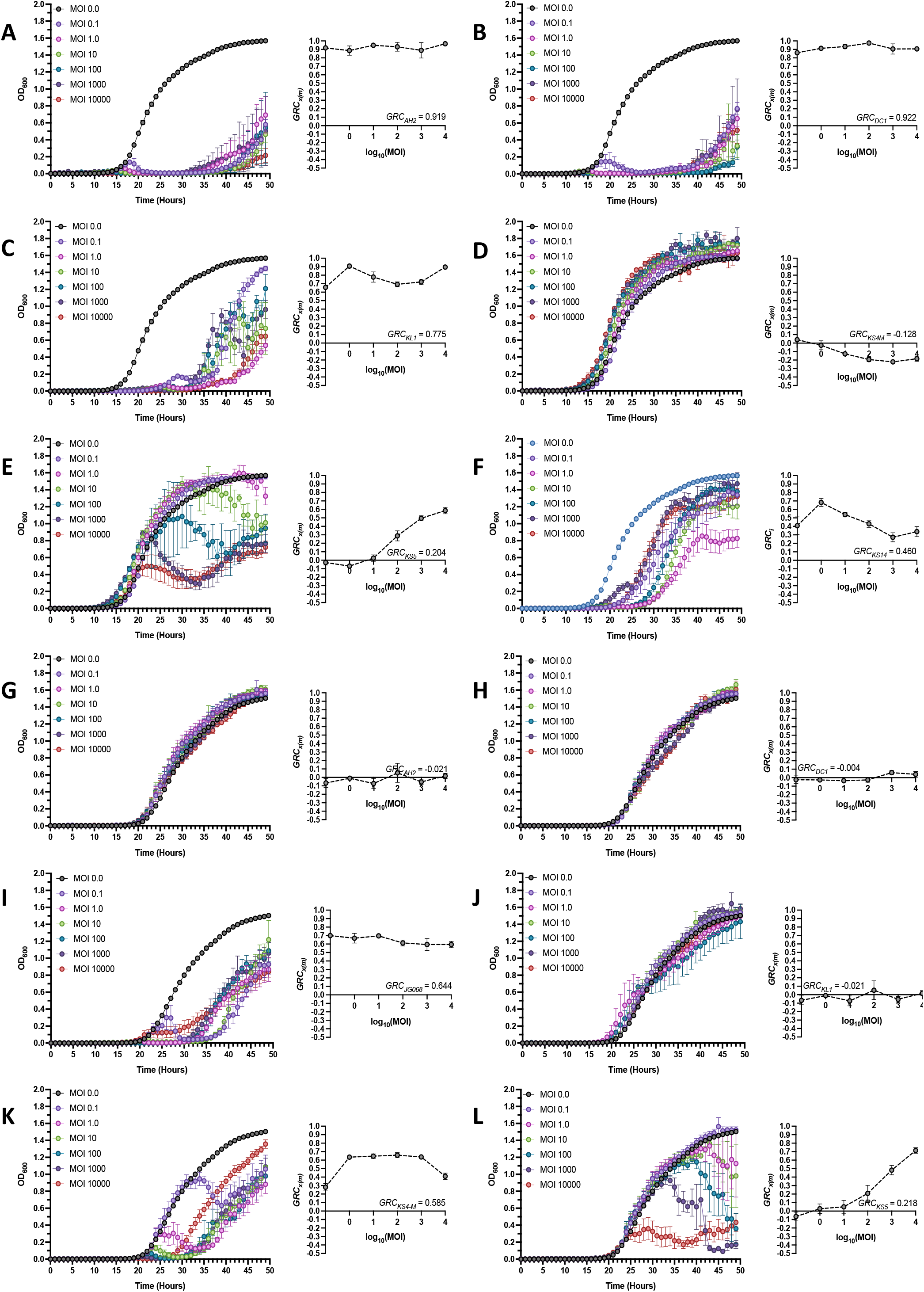
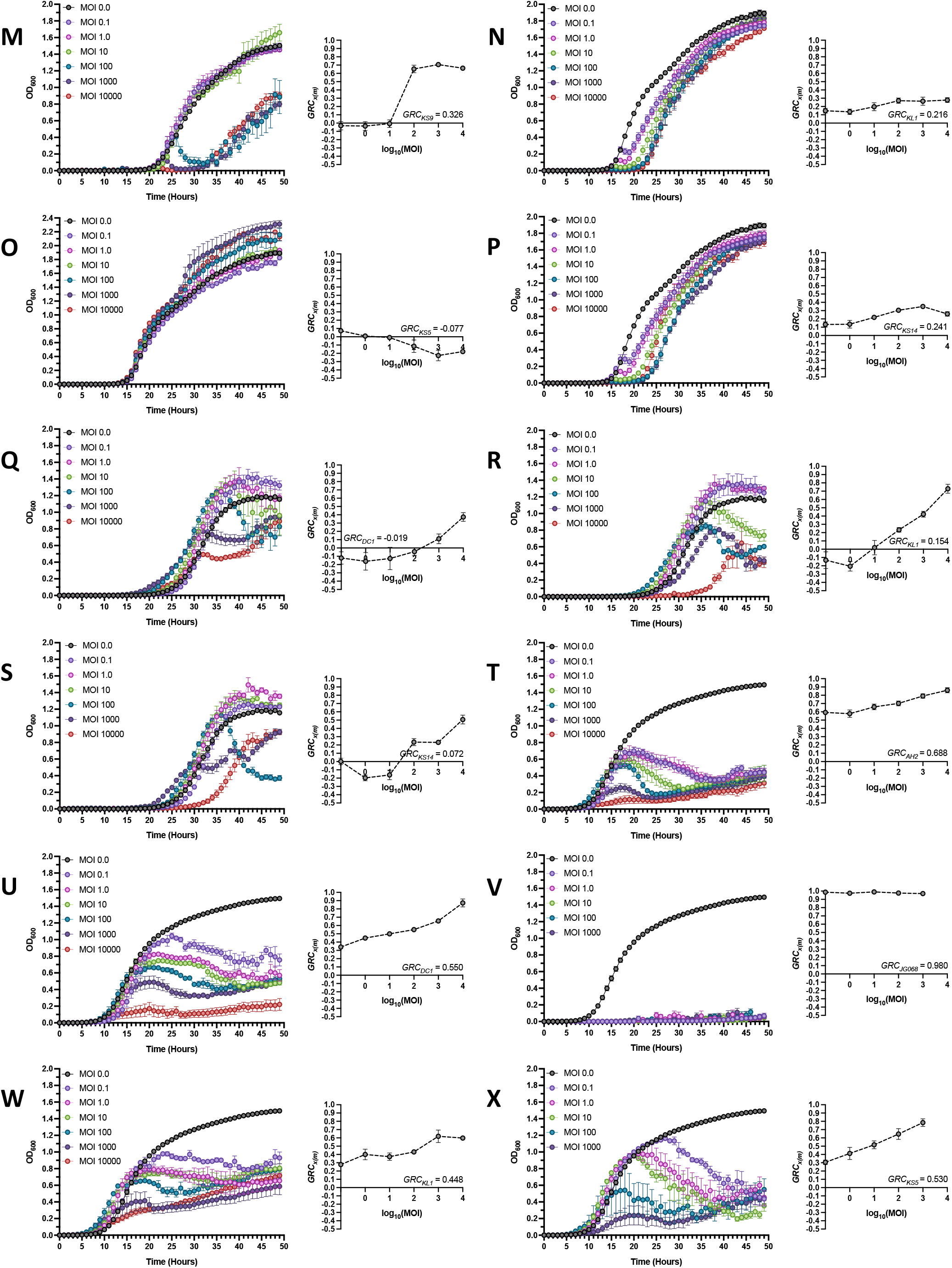
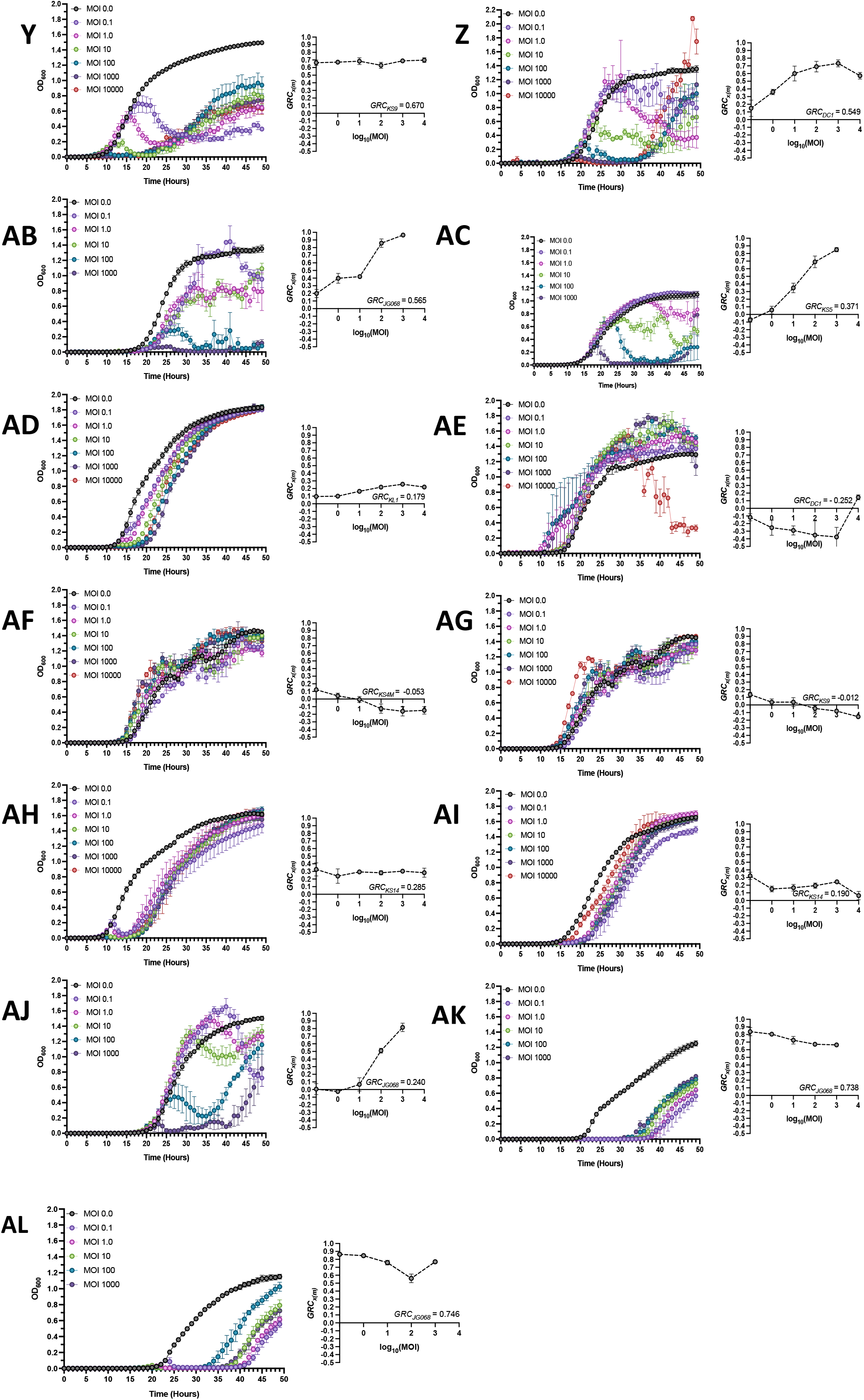
Growth Reduction Trends of Phages Targeting the Bcc. Bacterial growth curves (left) and associated Growth Reduction Coefficient (GRC) curves (right) of Bcc phages on *Burkholderia cenocepacia* strains **C6433** [phages AH2(c) (**A**), DC1 (**B**), KL1(c) (**C**), KS4M(c) (**D**), KS5(c) (**E**), KS14 (**F**)], **K56-2** [phages AH2(k) (**G**), DC1 (**H**), JG068 (**I**), KL1(k) (**J**), KS4M(k) (**K**), KS5(k) (**L**), KS9 (**M**)], **PC715J** [phages KL1(c) (**N**), KS5(c) (**O**), KS14 (**P**)], **AU41264** [phages DC1 (**Q**), KL1(c) (**R**), KS14 (**S**)], **AU41386** [phages AH2(c) (**T**), DC1 (**U**), JG068 (**V**), KL1(c) (**W**), KS5(k) (**X**), KS9 (**Y**)], **PC184** [phages DC1 (**Z**), JG068 (**AB**)], **J2315** [phage KS5(k) (**AC**)], **K63-3** [phage KL1(c) (**AD**)], and **CEP511** [phage DC1 (**AE**)]; *Burkholderia ambifaria* strains **LMG 19467** [phages KS4M(k) (**AF**), KS9 (**AG**)], and **LMG 17828** [phage KS14 (**AH**)]; *Burkholderia multivorans* strain **C5393** [phage KS14 (**AI**); *Burkholderia stabilis* strain **LMG 14294** [phage JG068 (**AJ**)]; and *Burkholderia dolosa* strains **AU0158** [phage JG068 (**AK**)], and **CEP021 CEP** [phage JG068 (**AL**)]. All experiments were performed at standard conditions across the standard MOI range of 0.1 to 10000, except for those phage-host pairs depicted in graphs **V, X, AB, AC, AJ, AK, AL**, for which the maximum available MOI was 1000. Black lines represent bacterial growth without phage (MOI 0.0), while coloured lines represent each of the investigated MOIs. 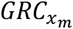 values were computed for each MOI using **equation 3**, and the *GRC*_*x*_ values for all phage-host pairs were then calculated using **equation 6** and are presented here for each pair. Note that the y-axes in graphs **O** and **Z** are slightly longer due to unusually high cell density resulting from phage treatment. Bars represent standard error of the mean of at least three biological replicates.

**Supplementary Figure 3:**
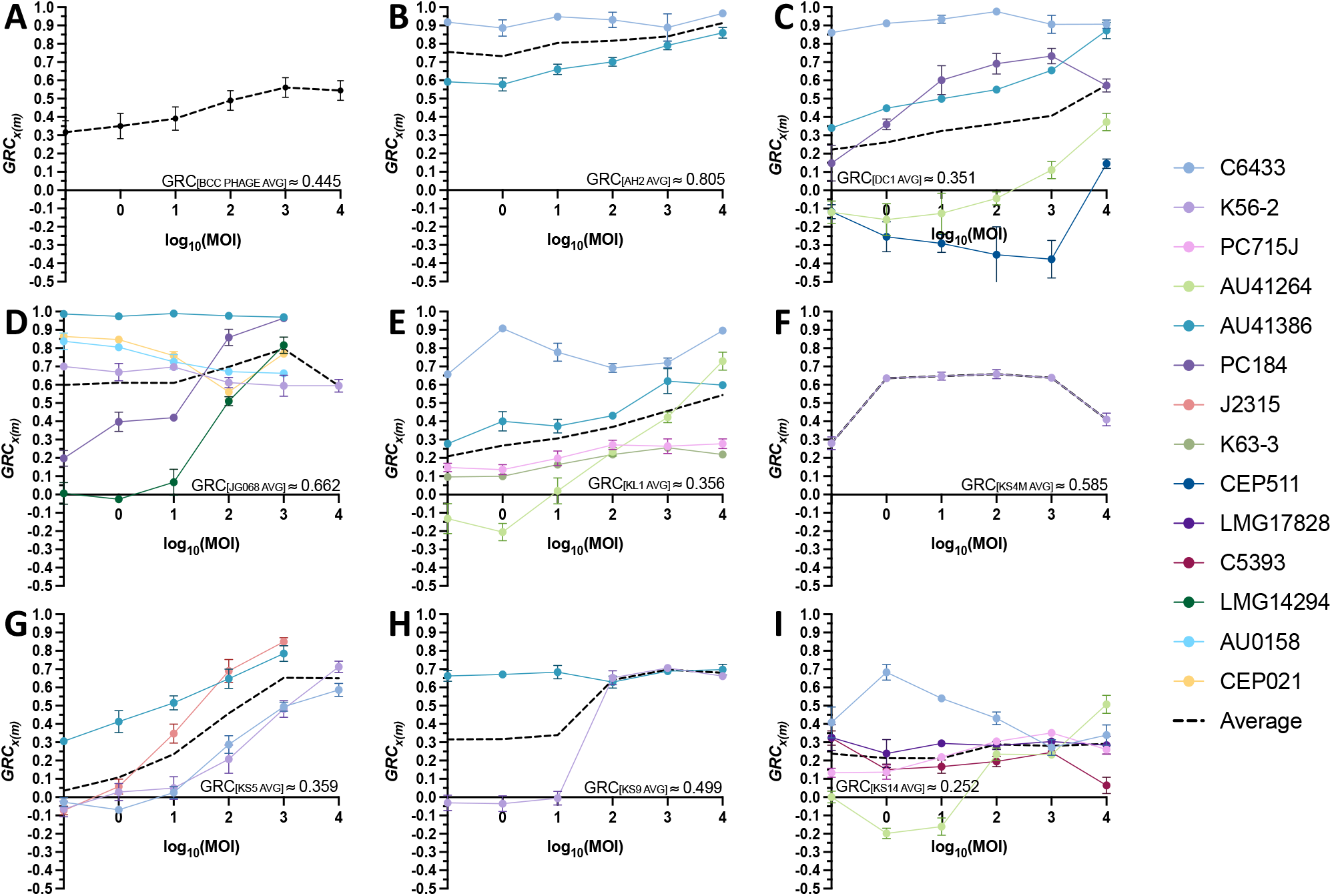
Mean 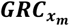 vs MOI trends of Bcc-targeting phages. Mean 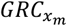 vs MOI trends and *GRC*_*x*_ values for a composite of all Bcc phages (**A**), as well as individual Bcc phages AH2 (**B**), DC1 (**C**), JG068 (**D**), KL1 (**E**), KS4M (**F**), KS5 (**G**), KS9 (**H**), and KS14 (**I**) on all tested host strains. For graphs depicting trends of individual phages (**B-I**), trends for each individual host strain are also shown to demonstrate the diversity of 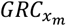 vs MOI trends even for individual phages. Bars represent standard error of the mean for at least three biological replicates.

**Supplementary Figure 4:**
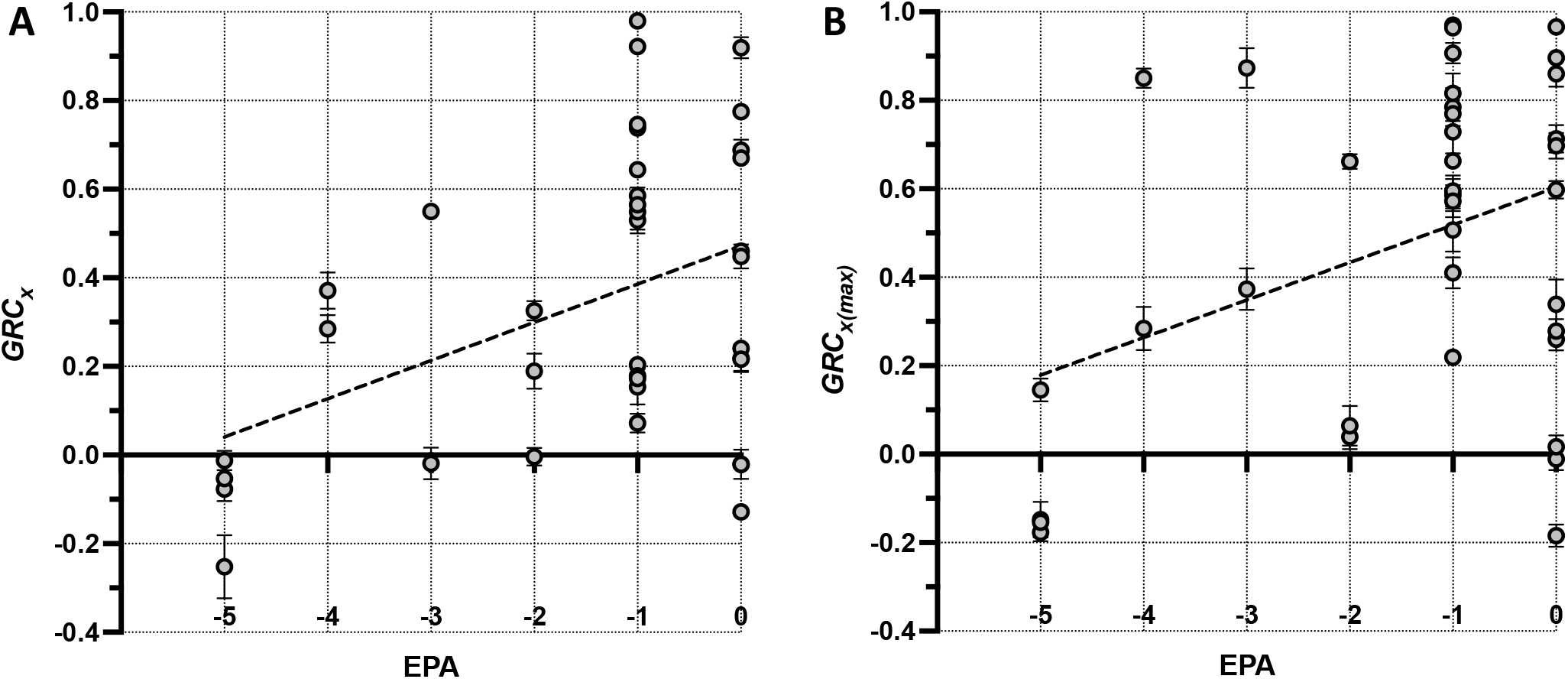
Correlation between Efficiency of Phage Activity (EPA) and the Growth Reduction Coefficient (GRC) of Phages Targeting the Bcc. EPA, *GRC*_*x*_, and 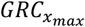 values of all investigated Bcc phage-host pairs, for infections conducted at standard conditions across the standard MOI range of 0.1 to 10000, were calculated using **equations 1, 6**, and **3**, respectively, and the EPA values were plotted against *GRC*_*x*_ (**A**) and 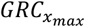 (**B**) values using x-y scatterplots. Dashed black lines represent lines-of-best-fit. Bars represent standard error of the mean for at least three biological replicates of GRC experiments.

**Supplementary Figure 5:**
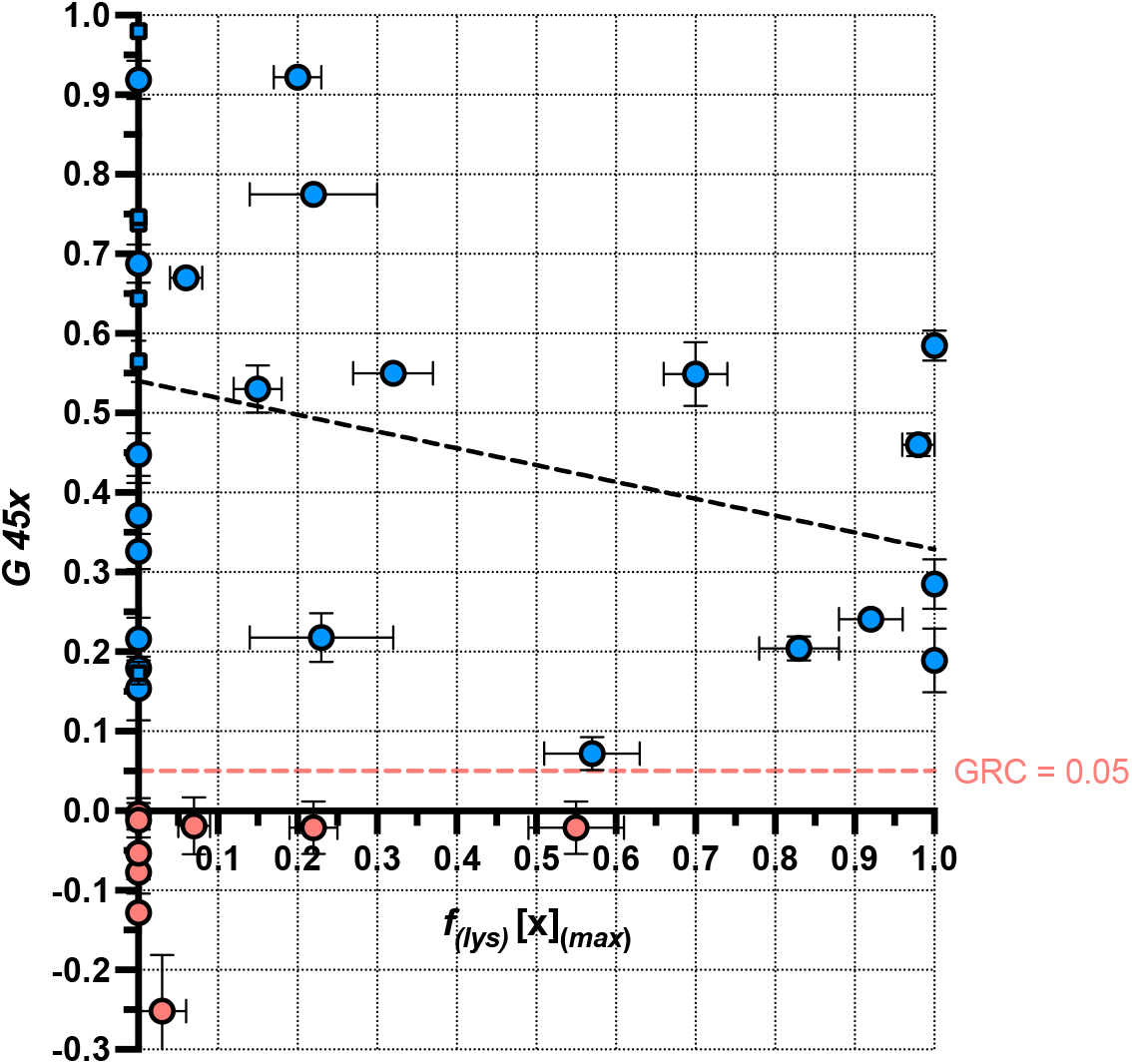
Lack of Significant Correlation between *GRC*_*x*_ and *f*_(*lys*)_ [*x*]_(*max*)._ *f*_(*lys*)_ values for infections conducted at the maximum available MOI, along with *GRC*_*x*_ values for infections conducted across the standard MOI range of 10^−1^ – 10^4^, all at standard conditions, were calculated for each investigated Bcc phage-host pair using **equations 9** and **6**, respectively, and were plotted using an x-y scatterplot. Points shown in red represent phage-host pairs which did not satisfy the GRC ≥ 0.05 criterion and thus fall below the red dashed line (representing a GRC of precisely 0.05), while points shown in blue represent phage-host pairs which did satisfy the GRC ≥ 0.05 criterion. Circles and squares represent phage-host pairs containing LC and OL phages, respectively. The black dashed line represents the line-of-best-fit for points shown in blue. Vertical and horizontal error bars represent standard error of the mean of at least three and four biological replicates, respectively.

**Supplementary Figure 6:**
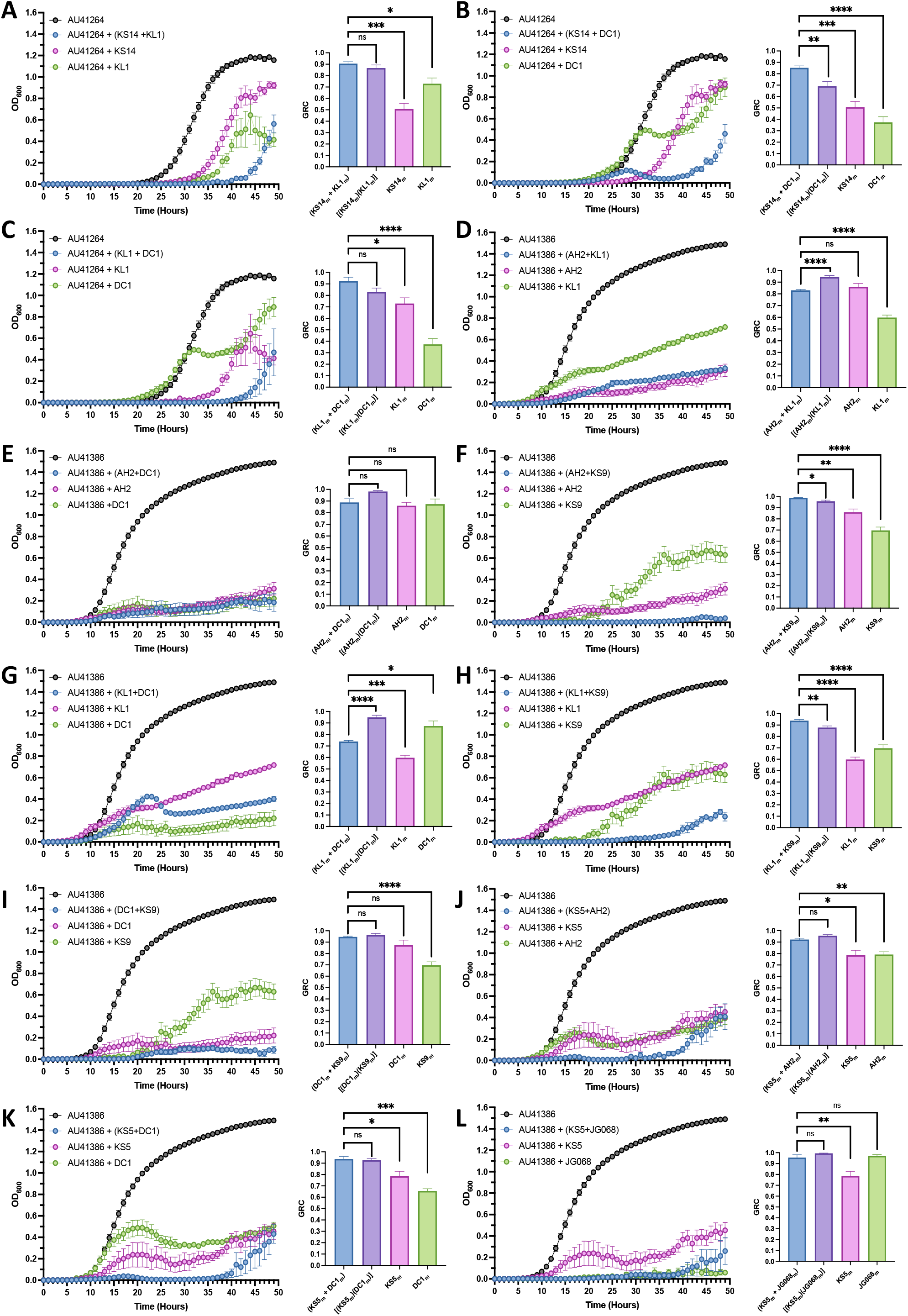
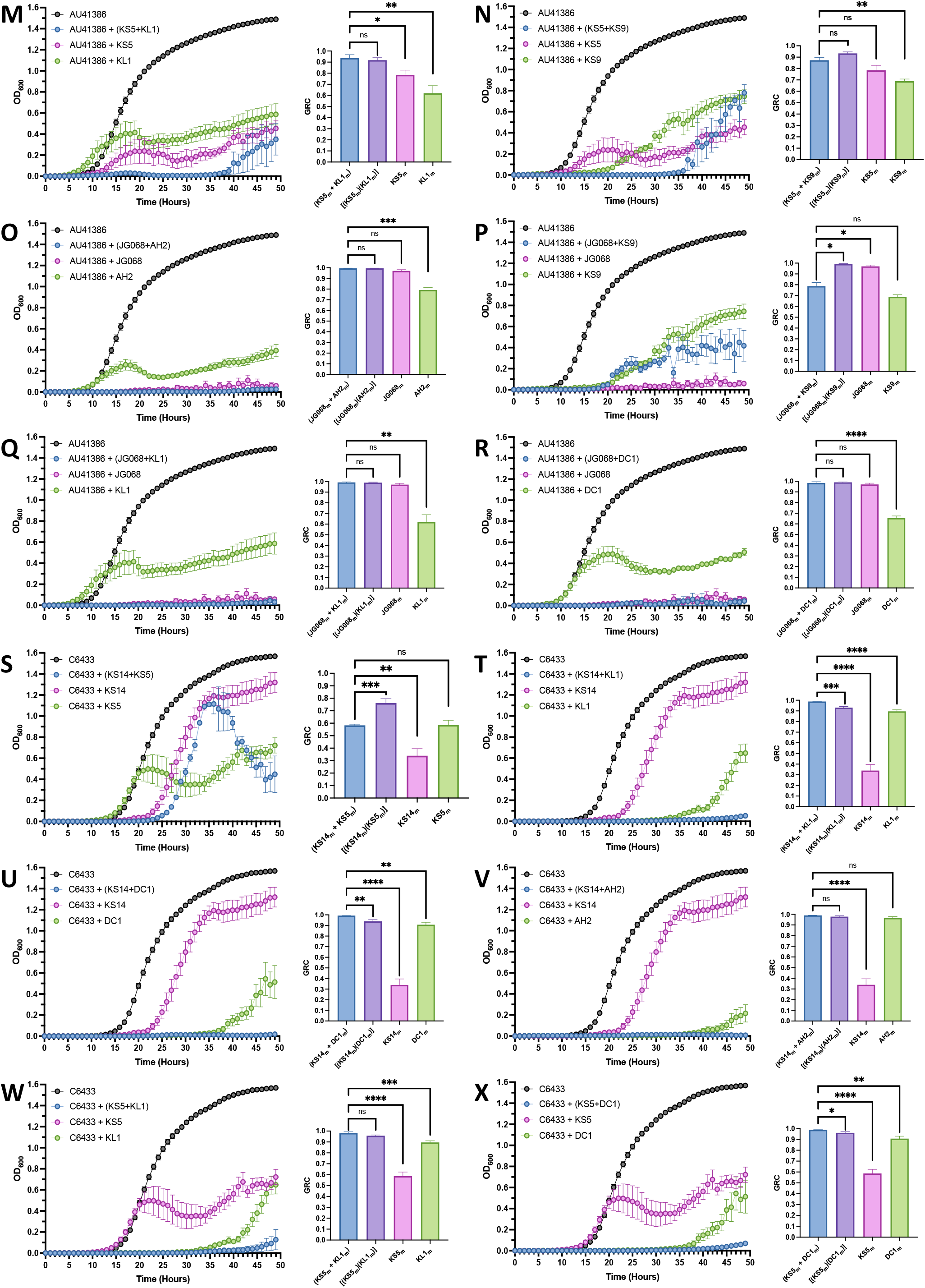
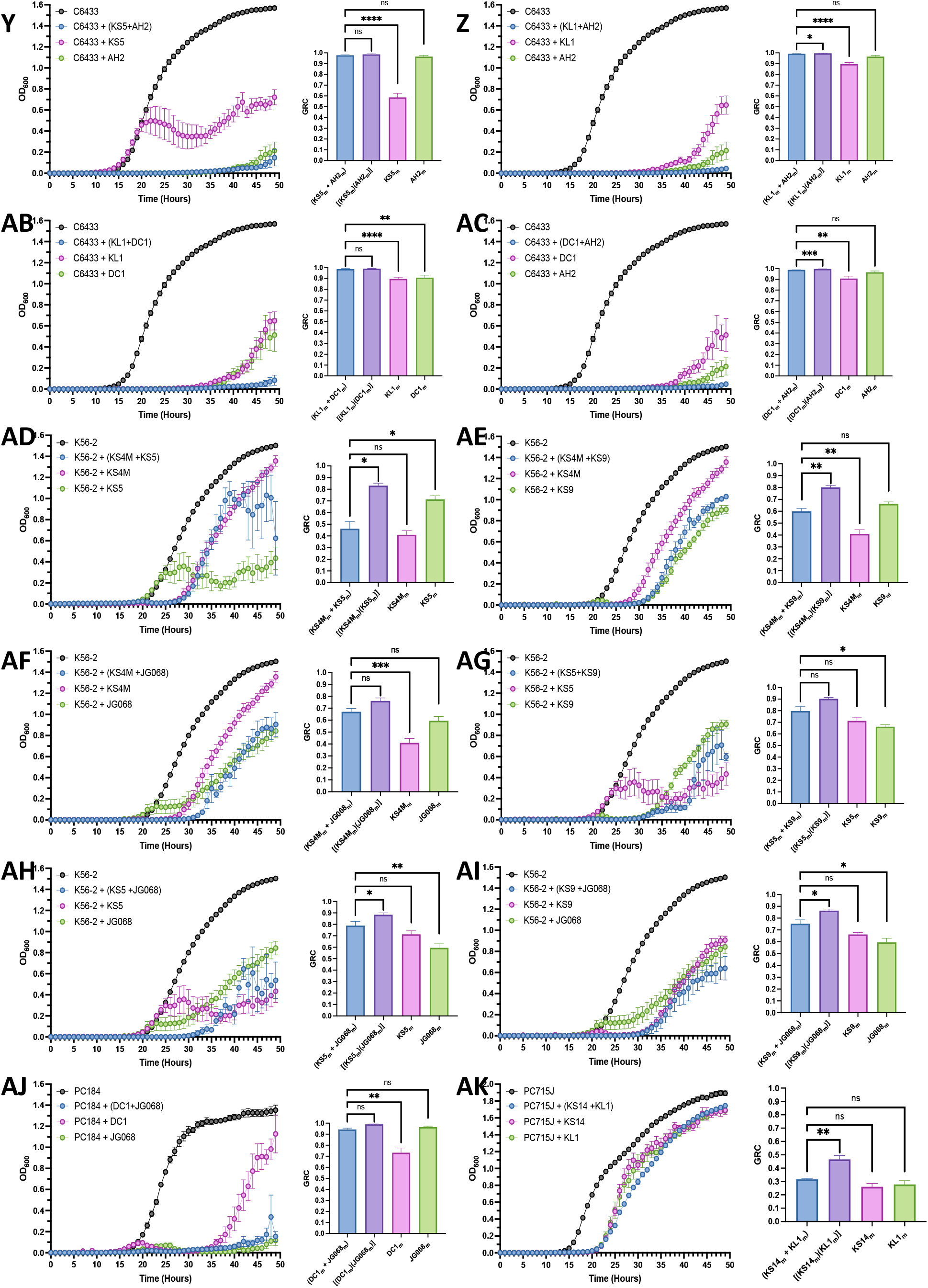
Interactions between Bcc-targeting Phages. Bacterial growth curves (left) and treatment GRC bar charts (right) for two-phage combinations on *Burkholderia cenocepacia* strains **AU41264** [KS14+KL1 (**A**), KS14+DC1 (**B**), KL1+DC1 (**C**)], **AU41386** [AH2+KL1 (**D**), AH2+DC1 (**E**), AH2+KS9 (**F**), KL1+DC1 (**G**), KL1+KS9 (**H**), DC1+KS9 (**I**), KS5+AH2 (**J**), KS5+DC1 (**K**), KS5+JG068 (**L**), KS5+KL1 (**M**), KS5+KS9 (**N**), JG068+AH2 (**O**), JG068+KS9 (**P**), JG068+KL1 (**Q**), JG068+DC1 (**R**)], **C6433** [(KS14+KS5 (**S**), KS14+KL1 (**T**), KS14+DC1 (**U**), KS14+AH2 (**V**), KS5+KL1 (**W**), KS5+DC1 (**X**), KS5+AH2 (**Y**), KL1+AH2 (**Z**), KL1+DC1 (**AB**), DC1+AH2 (**AC**)], **K56-2** [(KS4M+KS5 (**AD**), KS4M+KS9 (**AE**), KS4M+JG068 (**AF**), KS5+KS9 (**AG**), KS5+JG068 (**AH**), KS9+JG068 (**AI**)], **PC184** [DC1+JG068 (**AJ**)], and **PC715J** [KS14+KL1 (**AK**)]. Purple bars represent the product of the individual effectivenesses of the component phages, as computed using the right-hand side of **equation 11**. Black line represents bacterial growth without phage. Blue lines and bars represent bacterial growth with and GRC of the phage pair, while pink and green represent bacterial growth with and GRC of the individual phages. Bars represent standard error of the mean of at least three biological replicates. **** indicates p < 0.0001, *** indicates *p* < 0.001, ** indicates *p* < 0.01, * indicates *p* < 0.05, and ns indicates *p* > 0.05.

**Supplementary Figure 7:**
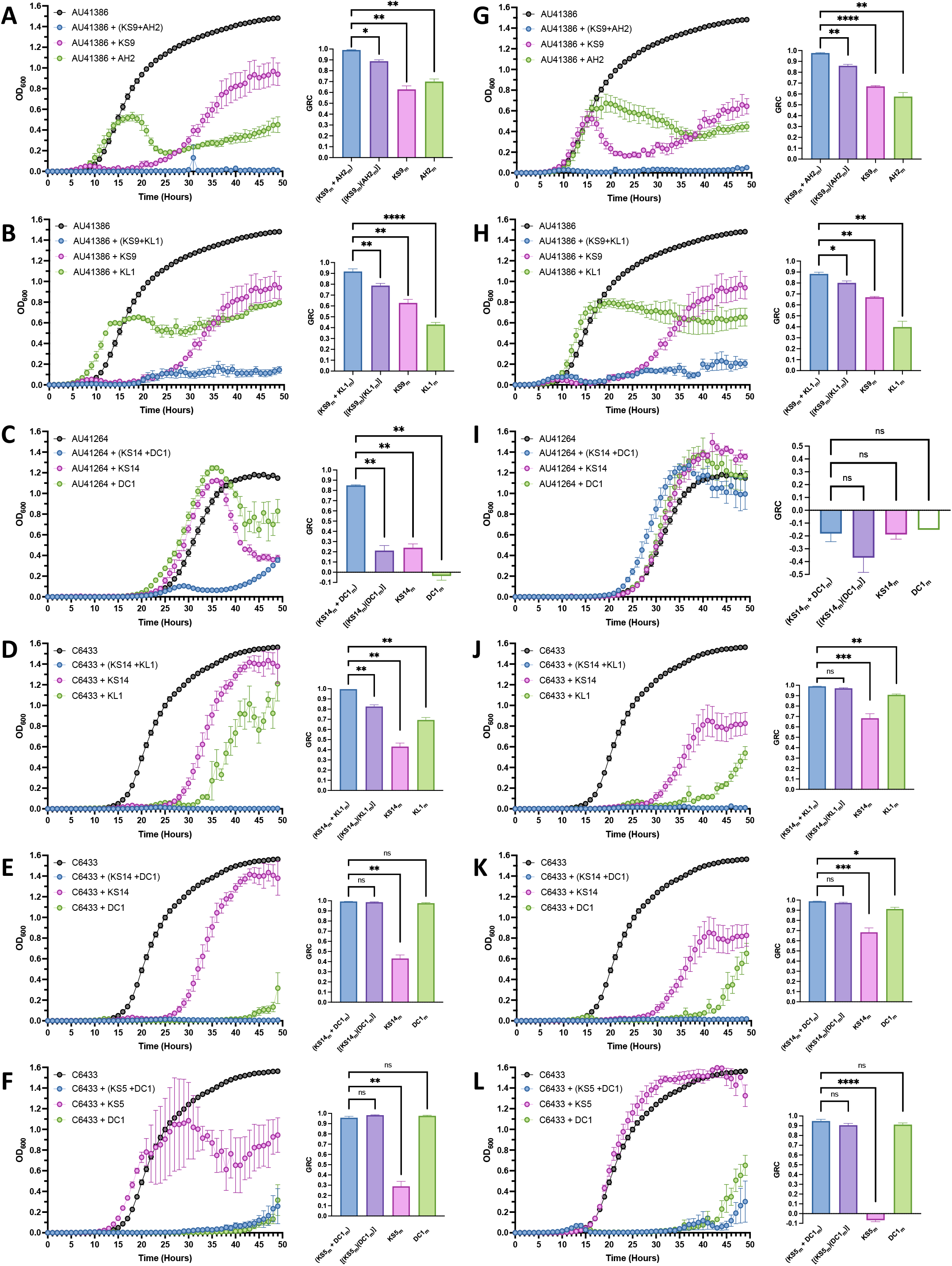
Interactions between Bcc-targeting Phages at lower MOIs. Bacterial growth curves (left) and treatment GRC bar charts (right) for the following phage combinations at MOIs 100 and 1, respectively: AH2 & KS9 (**A, G**) and KL1 & KS9 (**B, H**) on *Burkholderia cenocepacia* AU41386; KS14 & DC1 (**C, I**) on *Burkholderia cenocepacia* AU41264; KS14 & KL1 (**D, J**), KS14 & DC1 (**E, K**) and KS5 & DC1 (**F, L**) on *Burkholderia cenocepacia* C6433. Purple bars represent the product of the individual effectivenesses of the component phages, as computed using the right-hand side of **equation 11**. The black line represents bacterial growth without phage. Blue lines and bars represent bacterial growth with and GRC of the phage pair, while pink and green represent bacterial growth with and GRC of the individual phages. Bars represent standard error of the mean of at least three biological replicates. **** indicates *p* < 0.0001, *** indicates *p* < 0.001, ** indicates *p* < 0.01, * indicates *p* < 0.05, and ns indicates *p* > 0.05.

**SUPPLEMENTARY TABLE 1:** *Burkholderia cepacia* complex (Bcc) species & strains used in this study

| Species & strain | Source <sup>a</sup> | LD<sub>50</sub> in<br>*G. mellonella* | Bacterial<br>Inoculum | Phage<br>Inoculum | # phage<br>hits | # phages over<br>EPA threshold <sup>b</sup> | # phages over<br>GRC threshold <sup>c</sup> | # pairwise<br>combinations | Reference<br>or source |
| --- | --- | --- | --- | --- | --- | --- | --- | --- | --- |
| ***B. cenocepacia*** |  |  |  |  |  |  |  |  |  |
| C6433 | CF, Canada | 3.0 x 10<sup>4</sup> CFU | 10<sup>5</sup> CFU | 10<sup>4</sup> – 10<sup>9</sup> PFU | 6 | 6 | 5 | 10 | (40) |
| K56-2 | CF-e, Canada | 9.0 x 10<sup>2</sup> CFU | 10<sup>3</sup> CFU | 10<sup>2</sup> – 10<sup>7</sup> PFU | 7 | 7 | 4 | 6 | (40) |
| PC715J | CF, Canada | 4.0 x 10<sup>3</sup> CFU | 10<sup>5</sup> CFU | 10<sup>4</sup> – 10<sup>9</sup> PFU | 4 | 3 | 2 | 1 | (40) |
| AU41264 | CF, USA | No data | 10<sup>5</sup> CFU | 10<sup>4</sup> – 10<sup>9</sup> PFU | 3 | 3 | 3 | 3 | this study |
| AU41386 | CF, USA | No data | 10<sup>5</sup> CFU | 10<sup>4</sup> – 10<sup>9</sup> PFU | 6 | 6 | 6 | 15 | this study |
| PC184 | CF-e, USA | No data | 10<sup>5</sup> CFU | 10<sup>4</sup> – 10<sup>9</sup> PFU | 2 | 2 | 2 | 1 | (74) |
| J2315 | CF-e, UK | 1.0 x 10<sup>5</sup> CFU | 10<sup>5</sup> CFU | 10<sup>4</sup> – 10<sup>9</sup> PFU | 1 | 1 | 1 | 0 | (40) |
| K63-3 | CF, Canada | No data | 10<sup>5</sup> CFU | 10<sup>4</sup> – 10<sup>9</sup> PFU | 2 | 1 | 1 | 0 | (75) |
| CEP511 | CF, Australia | 8.0 x 10<sup>4</sup> CFU | 10<sup>5</sup> CFU | 10<sup>4</sup> – 10<sup>9</sup> PFU | 1 | 1 | 1 | 0 | (40) |
| ***B. ambifaria*** |  |  |  |  |  |  |  |  |  |
| LMG 19467 | CF, Australia | 8.0 x 10<sup>5</sup> CFU | 10<sup>5</sup> CFU | 10<sup>4</sup> – 10<sup>9</sup> PFU | 3 | 2 | 0 | 0 | (40) |
| LMG 17828 | Corn Roots | No data | 10<sup>5</sup> CFU | 10<sup>4</sup> – 10<sup>9</sup> PFU | 1 | 1 | 1 | 0 | (76) |
| ***B. multivorans*** |  |  |  |  |  |  |  |  |  |
| C5393 | CF, Canada | 3.0 x 10<sup>6</sup> CFU | 10<sup>5</sup> CFU | 10<sup>4</sup> – 10<sup>9</sup> PFU | 1 | 1 | 1 | 0 | (40) |
| C5274 | CF, Canada | 1.0 x 10<sup>6</sup> CFU | 10<sup>5</sup> CFU | 10<sup>4</sup> – 10<sup>9</sup> PFU | 1 | 0 | 0 | 0 | (40) |
| ***B. stabilis*** |  |  |  |  |  |  |  |  |  |
| LMG 14294 | CF, Belgium | 2.0 x 10<sup>6</sup> CFU | 10<sup>5</sup> CFU | 10<sup>4</sup> – 10<sup>9</sup> PFU | 1 | 1 | 1 | 0 | (40) |
| ***B. dolosa*** |  |  |  |  |  |  |  |  |  |
| LMG21443 | Rhizosphere | 4.0 x 10<sup>4</sup> CFU | 10<sup>5</sup> CFU | 10<sup>4</sup> – 10<sup>9</sup> PFU | 1 | 0 | 0 | 0 | (40) |
| AU0158 | CF, USA | No data | 10<sup>5</sup> CFU | 10<sup>4</sup> – 10<sup>9</sup> PFU | 1 | 1 | 1 | 0 | (77) |
| CEP021 | CF, USA | No data | 10<sup>5</sup> CFU | 10<sup>4</sup> – 10<sup>9</sup> PFU | 1 | 1 | 1 | 0 | (76) |
| **Total:** |  |  |  |  | **42** | **37** | **30** | **36** |  |

<sup>a</sup> Abbreviations: CF, cystic fibrosis infection; CF-e, strain that has spread epidemically among CF patients.
<sup>b</sup> In this study, the EPA threshold is arbitrarily set to EPA ≥ -5
<sup>c</sup> In this study, the GRC threshold is arbitrarily set to GRC ≥ 0.05

**SUPPLEMENTARY TABLE 2:** *Burkholderia cepacia* complex (Bcc) – targeting Bacteriophages used in this study

| Bacteriophage | Morphology | Source | Receptor | Lifestyle | Host Range [#Hosts]<sup>a</sup> | Reference |
| --- | --- | --- | --- | --- | --- | --- |
| AH2 | *Siphoviridae* | *Nandina* sp. soil | Unknown, not LPS | Lysogeny-capable | *B. cenocepacia* (C6433, K56-2, PC715J, AU41386, K63-3) [5 hosts] | (8, 42) |
| DC1 | *Podoviridae* | *Dracaena* sp. soil | Unknown | Lysogeny-capable; integrates | *B. cenocepacia* (C6433, K56-2, AU41264, AU41386, PC184, CEP511) [6 hosts] | (8, 43) |
| JG068 | *Podoviridae* | Sewage | LPS inner core | Obligately lytic | *B. cenocepacia* (K56-2, AU41386, PC184), *B. stabilis* (LMG 14294), *B. dolosa* (AU0158, CEP021). [6 hosts] | (8, 36) |
| KL1 | *Siphoviridae* | Sewage | Unknown, not LPS | Lysogeny-capable | *B. cenocepacia* (C6433, K56-2, PC715J, AU41264, AU41386, K63-3) [6 hosts] | (8, 42) |
| KS4M | *Myoviridae* | Mutant of DK4/BcepMu/KS4 | Unknown | Lysogeny-capable | *B. cenocepacia* (C6433, K56-2) *B. ambifaria* (LMG 19467) [3 hosts] | (8, 41) |
| KS5 | *Myoviridae* | Onion rhizosphere | LPS (lipid A proximal element) | Lysogeny-capable; integrates at 3' end of AMP nucleosidase | *B. cenocepacia* (C6433, K56-2, PC715J, AU41386, J2315), *B. ambifaria* (LMG 19467) [6 hosts] | (8, 41) |
| KS9 | *Siphoviridae* | Lysogen of *B. pyrrocinia* LMG 21824 | LPS (lipid A distal element) | Lysogeny-capable; integrates into 3' end of GTP cyclohydrolase II gene | *B. cenocepacia* (K56-2, AU41386), *B. ambifaria* (LMG 19467) [3 hosts] | (8, 41) |
| KS14 | *Myoviridae* | *Dracaena* sp. soil | Unknown | Lysogeny-capable; forms phagemid | *B. cenocepacia* (C6433, PC715J, AU41264), *B. ambifaria* (LMG 17828), *B. multivorans* (C5393, C5274), *B. dolosa* (LMG 21443) [7 hosts] | (8, 12) |

<sup>a</sup>These are the host ranges reported *in this study*, and differ from the host ranges reported when the phages were first isolated, which are listed comprehensively in (8). A bacterial strain was considered a viable host for a given phage if the phage-host pair yielded an EPA value $\geq -7$ .

**SUPPLEMENTARY TABLE 3:** Goodness-of-Fit and Significance Values for Figure 5A and Supplementary Figures 4 & 5

| Figure | Analysis | Goodness of Fit | Significance <sup>a</sup> |
| --- | --- | --- | --- |
| **Figure 5A:**<br>Correlation between $GRC_{x_m}$ and $f_{(lys)}$ at the maximum available MOI | Linear regression (all data points) | $R^2 = 0.03$ | $p > 0.05$ (ns) |
| | Linear regression (above GRC threshold) | $R^2 = 0.27$ | $p = 0.0033$ (**) |
| | Linear regression (above GRC threshold, excluding outlier group) | $R^2 = 0.67$ | $p < 0.0001$ (****) |
| | Linear regression (below GRC threshold) | $R^2 = 0.28$ | $p > 0.05$ (ns) |
| **Supplementary Figure 4A:**<br>Correlation between EPA and $GRC_x$ | Linear regression (all data points) | $R^2 = 0.18$ | $p = 0.0083$ (**) |
| **Supplementary Figure 4B:**<br>Correlation between EPA and $GRC_{x_{max}}$ | Linear regression (all data points) | $R^2 = 0.14$ | $p = 0.0211$ (*) |
| **Supplementary Figure 5:**<br>Lack of Significant Correlation between $GRC_x$ and $f_{(lys)}[x]_{(max)}$ | Linear regression (all data points) | $R^2 = 0.001$ | $p > 0.05$ (ns) |
| | Linear regression (above GRC threshold) | $R^2 = 0.10$ | $p > 0.05$ (ns) |
| | Linear regression (below GRC threshold) | $R^2 = 0.07$ | $p > 0.05$ (ns) |

<sup>a</sup> Significance refers to whether or not the slope of the line-of-best-fit is significantly different from zero.

